# Identification of letters distorted by physiologically-inspired spatial scrambling

**DOI:** 10.1101/2024.03.27.583720

**Authors:** Raffles Xingqi Zhu, Alex S. Baldwin

## Abstract

In the geniculostriate pathway of the human visual system, neuronal projections carry signals from a particular retinal locus in parallel from one anatomical area to the next. Imprecision in the fidelity of these projections would place constraints on the ability of the system to perform tasks requiring positional information. We investigated the impact that “spatial scrambling” between stages would have on visual performance. We consider two stages in a simple canonical model of the early visual cortex where scrambling might occur: either the input to the first orientation-tuned mechanisms (analogous to V1 simple cells), or the output from those mechanisms. These are referred as “subcortical” (SCS) and “cortical scrambling” (CS). We developed a wavelet decomposition and resynthesis algorithm to mimic these effects, and measured human performance in letter identification affected by the two types of scrambling. Our results showed SCS and CS have distinguishable effects on both perceived noisiness of letters and letter identification threshold. Comparing human performance against a suite of pre-trained and custom convolutional neural networks (CNNs) that were trained on the scrambled stimuli, relative efficiency (calculated from the ratio of human:CNN thresholds) is higher for CS than SCS. However, in modelling human inefficiency by reducing the proportion of wavelets available to the CNNs, humans are *less* efficient in CS than SCS. These differences in efficiencies show humans are better at processing orientation redundant stimuli (CS) than orientation noisy stimuli (SCS). We hypothesize this reflects differences in integration properties at the input and output stages of simple cells in the cortex.

**Author Summary:** The brain makes sense of the input from our eyes through a system where features are extracted and combined in successive stages. Our study concerns the spatial fidelity of the connections between visual areas. Previous behavioural and physiological evidence has suggested a scrambling of neuronal projections is present in biological visual systems. In our study, we investigate the ability of the human visual system to perform letter identification with stimuli affected by different types of on-screen distortions. These distortions simulate internal scrambling occurring at two early stages in the visual hierarchy. We used convolutional neural network (CNN) models as a benchmark, against which we compared human performance to find human efficiency in handling the distortions. We found that the type of scrambling in which humans were determined to have greater “efficiency” (relative to the CNNs) depended on the analysis used. The threshold magnitude of scrambling at which the letters could no longer be identified was greater for letters scrambled *after* the oriented features were extracted. Conversely, when efficiency was calculated by starving the CNNs of samples until their performance declined to the human level we instead found that the effective “number of samples” used by our humans was much higher for stimuli simulating scrambling *before* the oriented feature stage. These differences reflect how information is pooled and combined for these two stages.

## 1 Introduction

The rich external world we perceive through vision is the output of a hierarchical system, where representations are built up through multiple stages of visual processing (Hubel and Wiesel, 1962; DiCarlo et al., 2012; Riesenhuber and Poggio, 1999). Physiology puts limits on both the quantity and quality of information that gets transmitted at each stage. For example, information from the retina gets only partially transmitted to the cortex via the optic nerve (Kelly, 1962; Nirenberg et al., 2001) . Similar bottlenecks also exist in cortical processing, starting as early as V1. These are determined by the number of neurons, their energy consumption, and noise in the system (Li, 2014). A dominant topic within vision science has been the question of *how* the brain decodes a noisy input into an internal representation that drives our behaviour. To this end, it is argued that the brain performs attentional selection in the form of a saliency map (Li, 2002). to efficiently encode information (Clark, 2013; Rao and Ballard, 1999) (Barlow, 1956)

In addition to being hierarchical, the visual system is also topographical with visual maps corresponding to the outside world (Luo and Flanagan, 2007; Chklovskii and Koulakov, 2004; Gattass et al., 1981, 1988; Dumoulin and Wandell, 2008; Weinberg, 1997; Kaas, 1997; Groen et al., 2022). Deviations in the spatial organisation of projections between regions can result in a loss of spatial precision in the encoding of a feature. For example, in higher cortical areas, encoded representations of more complex stimulus features may be *less* specific in the positions of the features they encode (Gattass et al., 1981, 1988; Dumoulin and Wandell, 2008; Weinberg, 1997; Kaas, 1997; Groen et al., 2022).

Physiological studies have shown scatter in neuronal projections (Nishimoto et al., 2006; Tao et al., 2012; Zhang et al., 2013; Ringach, 2004). In animal models of amblyopia, this scatter is worse leading to more heterogeneous receptive fields (Tao et al., 2014; Swindale and Mitchell, 1994; Bi et al., 2011). Studies using fMRI found enlarged population receptive fields in amblyopic humans (Clavagnier et al., 2015; Hussain et al., 2015; Szinte et al., 2024; Li et al., 2007). A long history of behavioral studies have investigated the perceptual consequences of proposed intrinsic “scrambling” and “positional uncertainty” in visual tasks. These tasks investigated the effect of position noise on detection/localization (Michel and Geisler, 2011; Hess and Field, 1993, 1994; Morgan et al., 2012), bisection (Levi et al., 1987; Levi and Klein, 1986), shape//form discrimination (Watt and Hess, 1987; Keeble and Hess, 1999; Levi et al., 2000, 1999, 1997), contour integration (Hess et al., 1997; Baldwin et al., 2017),, and discrimination of natural images (Christensen et al., 2019). In amblyopia, it has been shown that the amblyopic eye’s percept is affected by a subjective “distortion” (Hess et al., 1978, 1990; Sireteanu et al., 1993; Barrett et al., 2003; Maruya et al., 2025). In normal vision, metameric stimuli have been developed that are indistinguishable when shown in the periphery. This phenomenon is thought to reflect a loss of spatial precision in the summary statistic encoding of features at greater eccentricities (Freeman and Simoncelli, 2011; .Broderick et al., 2023; Balas et al., 2009).

One approach for characterising internal scrambling or positional uncertainty of a system is the equivalent noise paradigm (Pelli and Farell, 1999; Pelli, 1981). These experiments are most typically conducted in the contrast domain by introducing task-relevant “external” noise to a stimulus and measuring thresholds in a range of different external noise levels: a “noise-masking function”. Models fit to these data quantify the external noise magnitude required to exceed the limiting effect of the system’s “internal noise” on performance (Pelli and Farell, 1999; Pelli, 1981). Separate from internal noise, efficiency (as a ratio between 0% to 100%) measures the gap in performance between humans and the theoretical ideal observer (a benchmark for the best performance that can be achieved on a particular task; Geisler, 2004). Comparing measurements of performance in high levels of external noise between humans and ideal observer also allows for direct measurement of efficiency (Pelli et al., 2006). These methods have revealed insights into how humans link contours (Hess and Demanins, 1998; Hess et al., 1997; Baldwin et al., 2017), integrate motion direction (Dakin et al., 2005; Pavan et al., 2023), and extracting orientation (Dakin, 2001).

In the current study, we want to investigate how internal scrambling may limit human efficiency in letter identification. For the loci of internal scrambling, we chose to focus on the geniculostriate pathway as it is arguably the most understood and the route through which visual information is transmitted from the retina to primary visual cortex (V1) via the lateral geniculate nucleus (LGN) (Hubel and Wiesel, 1962; Alonso et al., 2001). **Fig 1A** shows the classic model of oriented simple cell receptive field formation, pooling from subunits representing isotropically-tuned LGN afferents (Hubel and Wiesel, 1962; Hirsch et al., 1998). Here, we investigated the extent to which the simulated effects of scrambling could be differentiated when applied at two stages in this simple feed-forward hierarchical model. We developed an algorithm that decomposes a stimulus into a set of wavelets. To mimic internal scrambling, the stimulus can then be reconstructed either veridically, or with scrambling that simulates either: i) random jitter of the spatial location of oriented receptive fields (analogous to simple cells in V1), which we label “cortical scrambling” or CS (**Fig 1B**), or ii) random jitter of the location of the isotropic subunits which form those oriented receptive fields (analogous to LGN afferents) which we label “subcortical scrambling” or SCS (**Fig 1C**). Finally, we also investigated performance for non-scrambled stimuli presented in bandpass noise (BN). This allows us to connect our findings to more traditional contrast noise-masking approaches (Majaj et al., 2002; Pelli et al., 2004, 2006). **Fig 2** shows our BN, CS, and SCS stimuli for a range of noise and scrambling levels.

**Fig 1.**
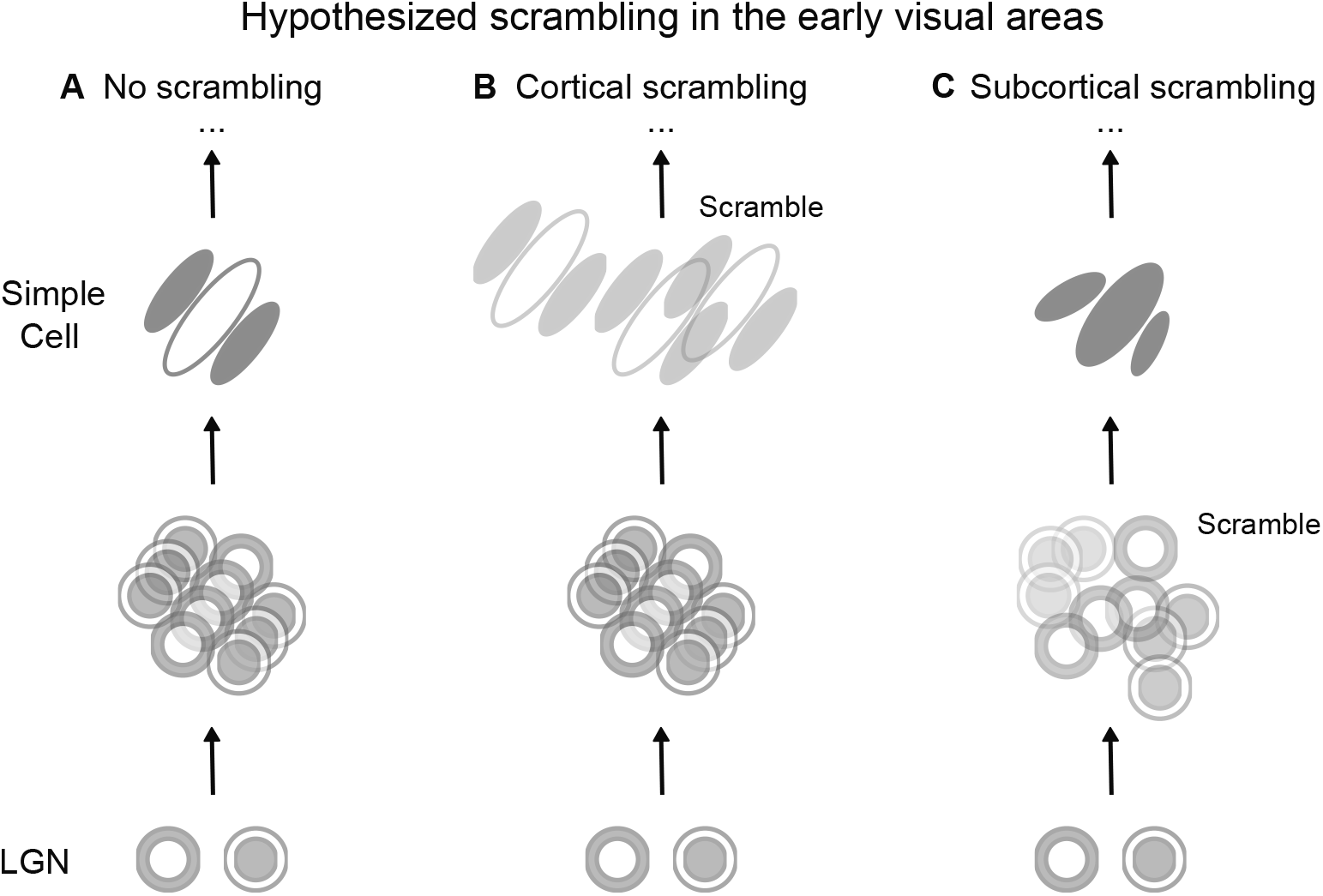
Hypothesized cortical vs. subcortical scrambling: Panel (**A**) shows a canonical view of V1 simple cell receptive field formation, being composed of inputs from the lateral geniculate nucleus (LGN) subunits. Panel (**B**) demonstrates the impact of scrambling simple cell projections to other cortical regions (“cortical scrambling”, or **CS**). Panel (**C**) shows the impact of scrambling the projections received from LGN subunits (“subcortical scrambling”, or **SCS**).

**Fig 2.**
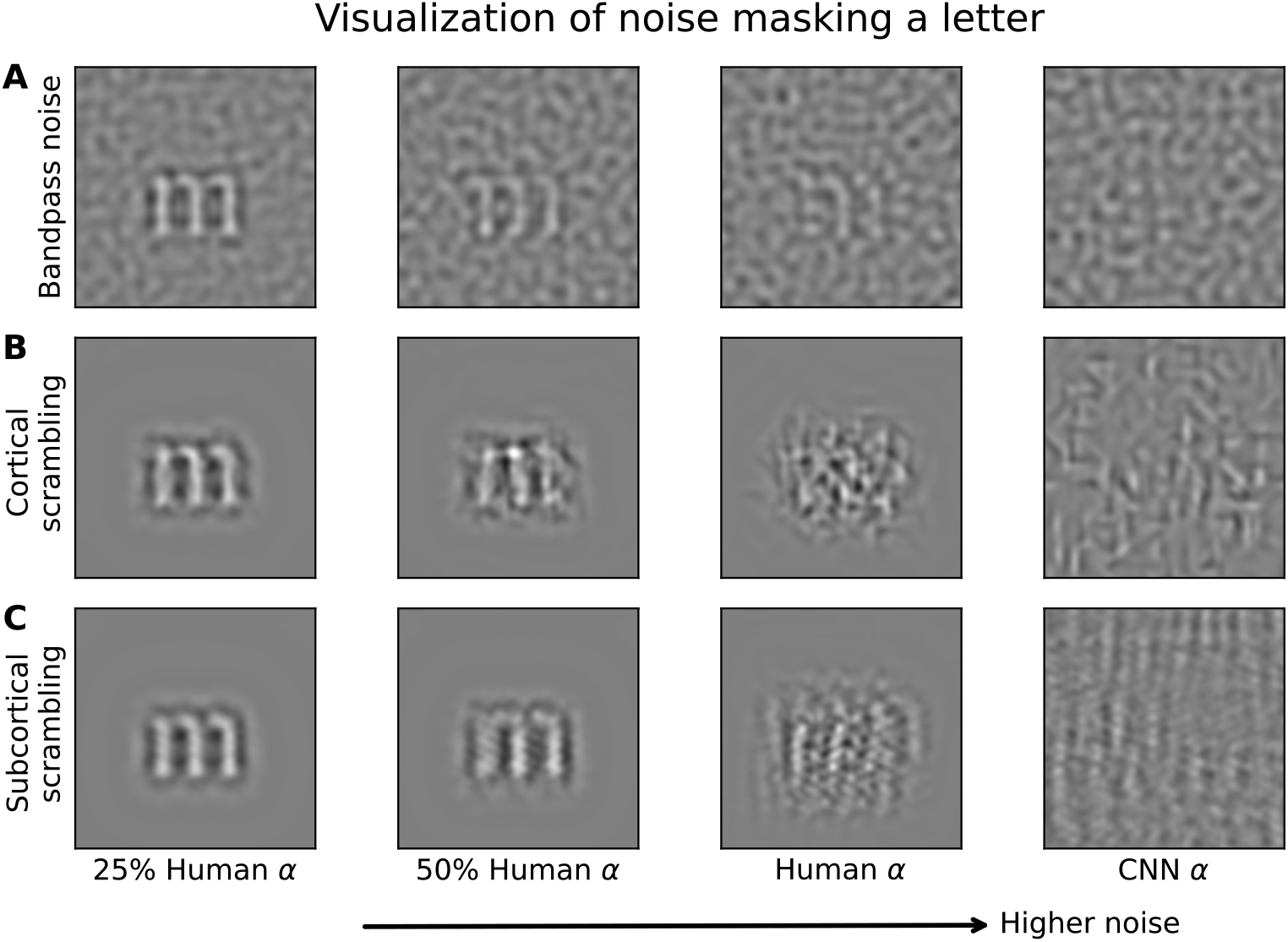
Visualizing the effect of masking noise used in this study. Spatial frequency of the letter was set to the optimal object spatial frequency for letter identification at 3 cycles/letter. The example uses a bandpass letter “m”, showing the effect of increasing noise magnitude from left to right for our three conditions: (**A**) bandpass noise (BN), (**B**) cortical scrambling (CS), and (**C**) subcortical scrambling (SCS). For illustration, noise levels are shown at fractions of human threshold (*α*), representing 62% correct in the letter identification task (**Fig 6**) for humans. Stimuli at *α* of convolutional neural networks (CNNs) (**Fig 5**) are also shown for reference. For BN (**A**), noise magnitude is quantified in the relative RMS contrast of the noise compared to the target. For CS (**B**) and SCS (**C**), noise magnitude is quantified as the standard deviation of the Gaussian from which position offsets of the wavelets or subunits are drawn.

Letter identification has clinical relevance in the measurement of letter acuity (Chung et al., 2002b; Pelli et al., 2004, 2006; Chung et al., 2002a),, a long-standing importance to the field of computer vision in Optical Character Recognition (Zhang et al., 2015; Bottou et al., 1994), and plays an important role in advancing theories and models of invariant object recognition (Grainger et al., 2008). Conveniently human psychophysics research has identified the optimal spatial frequency band (around 3 cycles/letter) that supports letter identification (Majaj et al., 2002; Chung et al., 2002b; Solomon and Pelli, 1994). This helped simplify our stimulus generation through presenting spatially-bandpass letters at that optimal spatial frequency, which should limit the contribution of complex cross-frequency gain control effects (Heeger, 1992; Bonds, 1989; Brouwer and Heeger, 2011; Eckstein et al., 1997; Foley, 1994; Meese and Holmes, 2006).

As letter identification is a classification task, this approach also facilitated the use of CNN models in our study. These are emerging as promising models against which human object recognition can be compared (Majaj and Pelli, 2018; Lindsay, 2021; Wichmann and Geirhos, 2023; Yamins and DiCarlo, 2016). It is difficult to develop an analytical ideal observer to predict the impact our CS and SCS manipulations would have on letter identification. Without this ideal observer, we also cannot present the efficiency for CS and SCS letter identification on an absolute scale. Instead, we chose to compare human performance against that of a range of CNN models (which, after Kanwisher et al., 2023, may be “optimised” but likely not “optimal”). To have networks optimised specifically for the task of interest, we trained a set of twenty networks from scratch following an architecture search (Francl and McDermott, 2022). We also investigated the behaviour of a varied set of networks pre-trained on ImageNet: AlexNet (standard feedforward CNN), ResNet50 (with skip connections), VGG19 (stacking small filters only), and CORnetS (with recurrent connections to mimic human vision) (Krizhevsky et al., 2012; He et al., 2015; Simonyan and Zisserman, 2015). Benchmarking human performance against that of CNNs allowed us to calculate relative efficiency for both CS and SCS on an objective scale.

To preview our results, humans exhibited higher relative efficiency for cortical scrambling (CS) than subcortical scrambling (SCS) (**Fig 6D-F**). Defining sampling efficiency as the proportion of wavelets made available to the CNNs to match human performance, we instead find more samples were needed in SCS than CS (**Fig 6G-I**). These results suggest difference in processing constraints between input and output stages of simple cells in the cortex.

## 2 Results

### 2.1 Experiment 1: Matching perceived scrambling magnitude

We performed a matching experiment to compare the subjective magnitude of perceived “scrambling” that resulted from our Cortical Scrambling (CS) and Subcortical Scrambling (SCS) algorithms. Although both kinds of scrambling were quantified by the standard deviation (*σ*) of the Gaussian process (in degrees of visual angle or “deg”) used to perturb the locations of the features, we nevertheless wished to determine whether one type might result in a greater subjective percept of “scrambling”. At the same time, we also sought to determine whether either type of scrambling might saturate, such that increases in standard deviation would have a diminishing effect on perceived scrambling beyond some critical magnitude. To answer these questions, we performed an experiment where participants matched the perceived scrambling of two letters affected by either CS (with scrambling magnitude *σ*CS) or SCS (with scrambling magnitude *σ*SCS).

We measured matches both between the same type of scrambling (matching CS to CS, or SCS to SCS) and between the two different types of scrambling (see **Methods** Section 4.7). In all comparisons, we found the matching relationship to be well-characterized by a simple linear fit on double-log axes (**Table 1**). When matching the scrambling magnitude of two letters affected by the same type of scrambling, a veridical match would have a slope of unity and an intercept of zero (*y* = *x*). We found participants were able to make matches close to this veridical relationship for both CS and SCS. This validates their ability to make accurate matches of perceived scrambling magnitude.

**Table 1.** Details of linear fits to scrambling matching experiment data on double-log axes. Cross-scrambling-type conditions are plotted in **Fig 3** Reduced refers to fitting one linear model to matching data in both directions.

| Condition | Parameters | Value | 95% CI | R<sup>2</sup> |
| --- | --- | --- | --- | --- |
| SCS reference<br>SCS match | Intercept | -0.02 | [-0.10 0.05] | 0.94 |
|  | Slope | 1.01 | [0.97 1.04] |  |
| SCS reference<br>CS match | Intercept | 0.02 | [-0.07 0.12] | 0.93 |
|  | Slope | 1.16 | [1.11 1.20] |  |
| SCS reference<br>CS match<br>(reduced) | Intercept | 0.02 | [-0.04 0.22] | 0.99 |
|  | Slope | 1.12 | [1.09 1.21] |  |
| CS reference<br>SCS match<br>(reduced) | Intercept | -0.02 | [-0.19 0.04] | 0.96 |
|  | Slope | 0.89 | [0.83 0.92] |  |
| CS reference<br>SCS match | Intercept | -0.26 | [-0.36 -0.17] | 0.87 |
|  | Slope | 0.81 | [0.76 0.85] |  |
| CS reference<br>CS match | Intercept | -0.08 | [-0.17 0.01] | 0.92 |
|  | Slope | 0.99 | [0.95 1.03] |  |

For matches between the two types of scrambling, the relationships were still linear (on our log-log axes, see Figure 3) however the slope and intercept values varied. We performed the match in both directions, adjusting scrambling magnitude of a CS letter to match a SCS letter (**Fig 3A**) and the other way around (**Fig 3B**). We found more SCS was needed to match the perceived noisiness of CS at lower scrambling levels. However, the perceived noisiness of the SCS increased more rapidly with the nominal noise level so that at the higher magnitudes tested the perceived “noisiness” of the two manipulations became similar for stimuli generated with the same *σ*.

**Fig 3.**
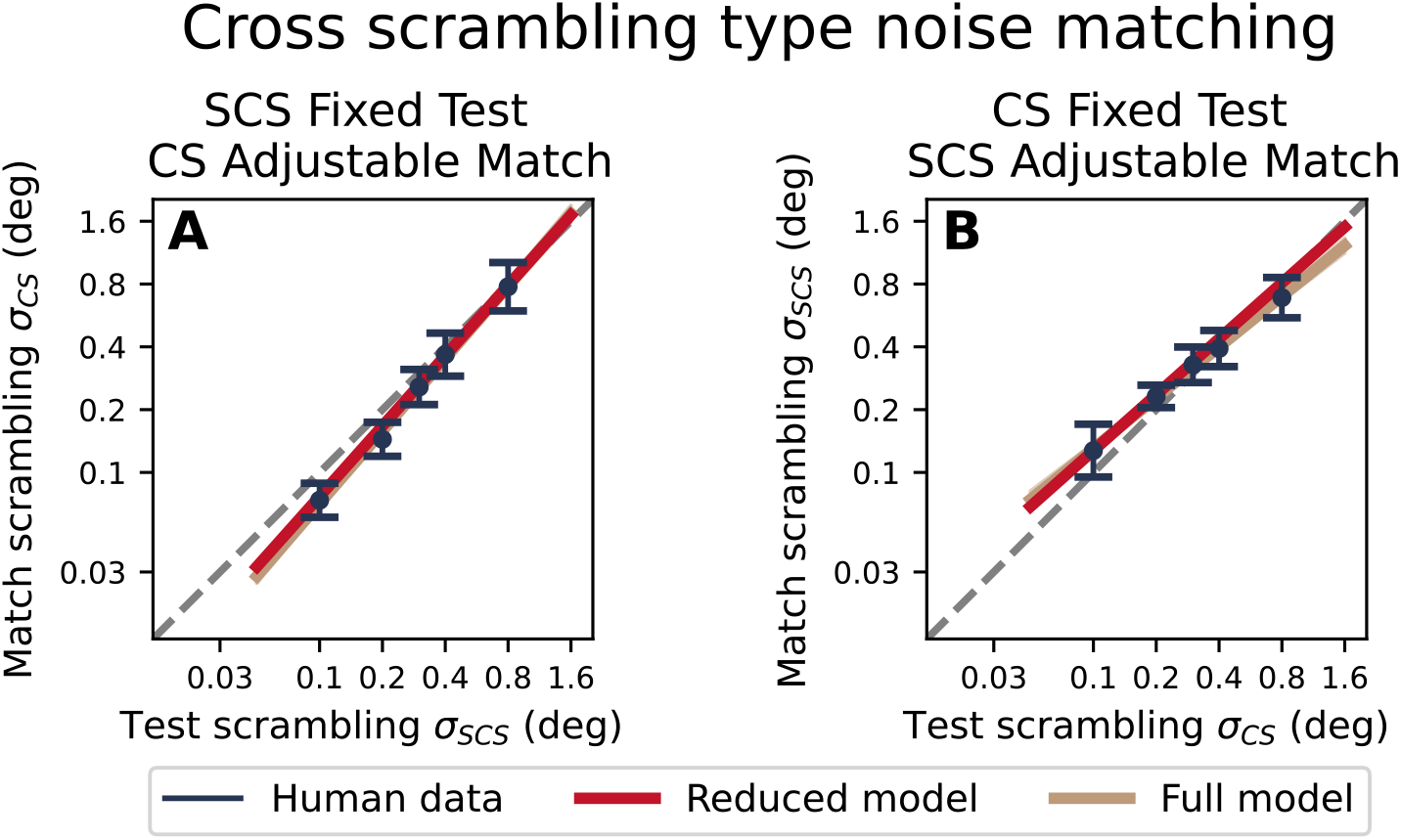
Matching cortical scrambling and subcortical scrambling (and vice versa). (**A**) shows results of matching a cortical scrambled (CS) letter (the match) to a subcortical scrambled (SCS) letter (the test). (**B**) shows the opposite match. The dashed grey line shows the unity line. The best-fit line to the data, fit separately for panels (**A**) and (**B**) is shown in gold. The best-fit line using the reduced model (common relation with a single set of slope and intercept) is shown in red. Markers show mean with standard deviation of the human data pooled across five participants.

We would expect that the equivalent SCS magnitude to match an CS stimulus could be predicted by the inverse of the rule for matching a CS stimulus to an SCS stimulus. We performed an analysis to determine whether the same rule could account for matches in both directions (CS to SCS and vice-versa) in a “reduced” model or if the two directions were better accounted for by two different linear relationships (the “full” model). A nested model hypothesis test found the full model to be superior (p < 0.001, nested model hypothesis test; details in **Table 1** of Supporting Information). Inspecting **Fig 3**, we see that the full model differs from the reduced model by better accounting for the condition where an SCS stimulus was matched to a reference CS stimulus of the highest scrambling magnitude tested. It is worth noting that the same stimulus magnitude was matched in the opposite direction with *σ*CS ≈ *σ*SCS so we conclude this deviation is likely an artefact of our testing procedure (for example, it may be that the linear matching relationship *does* break down for magnitudes beyond our chosen range). On that basis, and justified by the overall strong performance performance of our reduced model in accounting for all other data points, we propose that the two types of scrambling (within the range used in this study) follows the relationship:

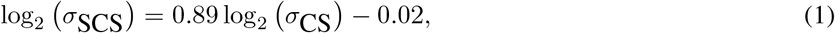

this is approximated by a simple power law where

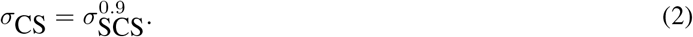

### 2.2 Experiment 2: Identification of letters affected by noise or scrambling

In our main experiment, we investigated how the two types of scrambling (CS and SCS) affected performance on a letter identification task, and how this compared to the identification of letters in bandpass noise (BN). We measured the magnitude of noise (BN) or scrambling (CS or SCS) that would reduce letter identification performance to our threshold level (62% correct). Example psychometric functions from a single participant are shown in **Fig 4A-C**, with the average thresholds calculated over 20 participants shown in **Fig 4D-F**. Separate thresholds were calculated for participants using their dominant (D) and non-dominant (ND) eyes. We found participants were significantly more resilient to SCS (higher threshold) using their D eye (t-test; *t*_19_=2.18, p=0.042). We did not find the same dominant eye advantage for BN (*t*_19_=-1.06, p=0.30) or CS (*t*_19_=0.05, p=0.96). These results are intriguing, as our SCS manipulation would correspond to type of scrambling we would expect to occur *before* binocular combination. To simplify the remainder of our analyses we will combine data from the two eyes of each participant by taking their average, though we will return to this point in the **Discussion**.

**Fig 4.**
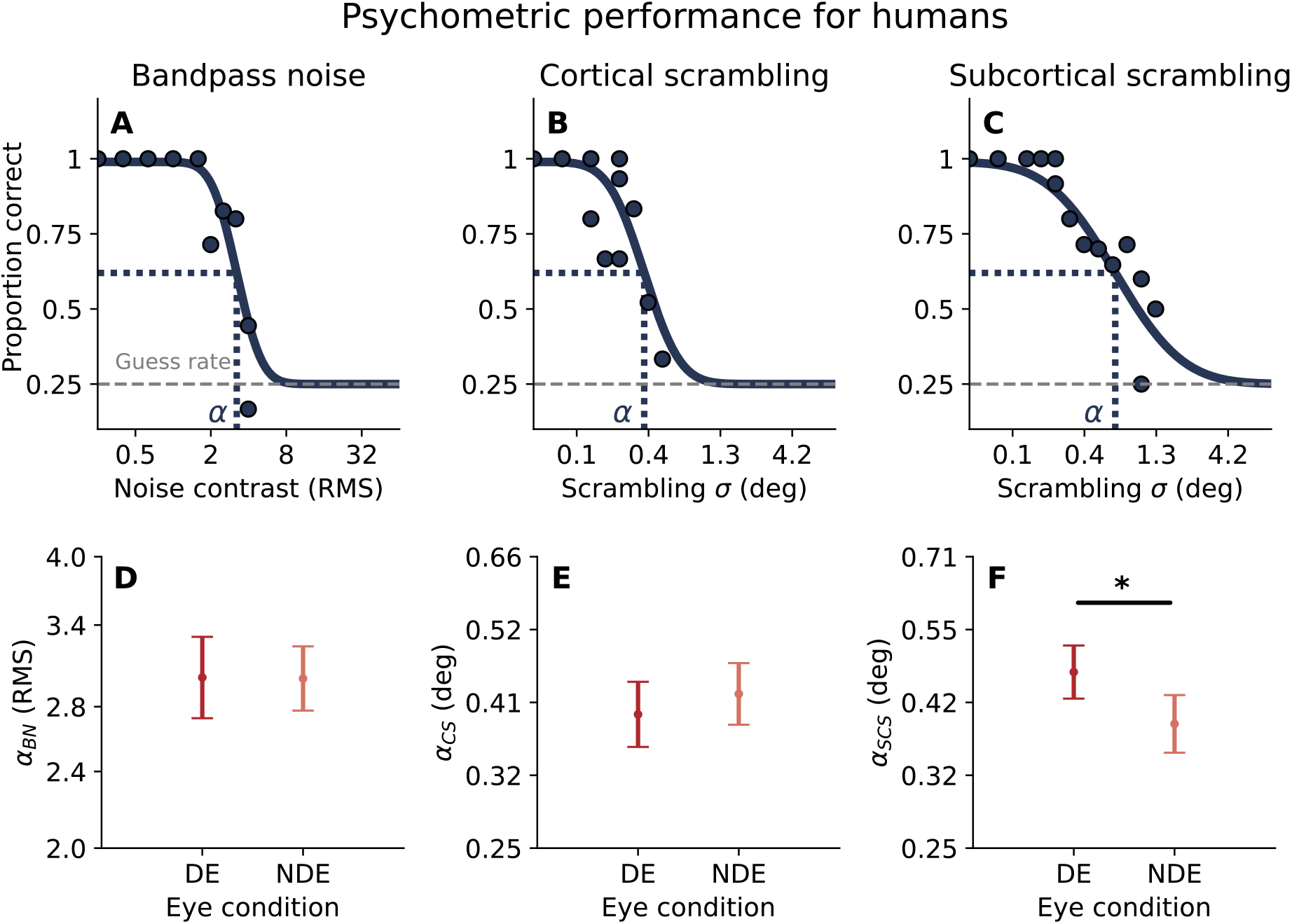
The top row (**A**-**C**) show example data from one human participant with the psychometric function fit by our maximum-likelihood method. The bottom row (**D**-**F**) shows mean noise / scrambling thresholds across 20 human subjects (with error bars showing 95% CI).

### 2.3 Performance of CNN models

The thresholds measured in our three conditions (BN, CS, and SCS) are not directly comparable, as the parameters controlling the magnitude of noise or scrambling are different in each condition. By finding the performance of CNN models optimised to perform the identification task under each stimulus condition, we sought to use them as a benchmark against which human performance could be compared. The relative efficiency, defined as the ratio of human performance to the performance of a model observer (Tanner and Birdsall, 1958; Pelli et al., 2006),allows for an objective comparison between different stimulus conditions. The models used in these comparisons are often “ideal observers” that make optimal use of all available information to perform the task (Geisler, 2004, 1989). For the two scrambling conditions tested in this study we did not have ideal observers, and so instead use our CNN models as a reference (while bearing in mind they likely fall short of the theoretical “best possible” performance).

We looked at the performance of both custom CNNs and pre-trained networks popular in the literature (that underwent transfer learning on our task). For our custom CNNs, to generate an optimised set for each noise condition we first performed an architecture search by generating 100 architectures with random hyper-parameters and picked the top twenty best-performing architectures (histograms of hyperparameters shown in **Fig 4** of Supporting Information). These twenty architectures were then trained from scratch until convergence. For the pre-trained networks, we chose VGG19, AlexNet, ResNet, CORnetS pre-trained on ImageNet. In our set of models, we also include a template matching (TM) model. This is the ideal observer model for identifying letters in white noise Van Trees (1968); Pelli et al. (2006); Ziskind et al. (2014). We therefore expect it to be optimal for the BN condition (and so would expect CNNs trained to perform the BN task to reach its performance level).

The psychometric functions for our computer models are shown in **Fig 5A-C**. The psychometric threshold *α* for various models are shown in **Fig 5D-F**. In each column, we see the best-performing custom CNNs for each stimulus condition were those that were *trained* on that same condition, i.e. the best-performing custom CNNs for the BN condition were those that were trained to identify BN stimuli. This is evidence that identification of BN stimuli is best supported by a distinct strategy. This is unsurprising, as the BN task (identification of a letter in-place without distortion) differs fundamentally from that of either of our scrambling conditions. We learn more from the comparison between the CS and SCS conditions. Although the generation of these stimuli involves distinct operations, it would nevertheless be possible that a similar strategy would be optimal to identify scrambled letters under either manipulation. Such an equivalence, had we found it, would limit our expectations for any ability to distinguish the two types of scrambling through behavioural measurements. Instead, we find CS-trained networks outperform SCS-trained networks when the task is to identify CS stimuli, and vice-versa. These differences were tested using a Mann-Whitney U test (p < 0.0001 for pairwise comparisons among BN CNN, CS CNN, and SCS CNN for all three noise conditions). These results echo previous works that have shown CNNs do not generalize well to out-of-distribution samples (Geirhos et al., 2018).

**Fig 5.**
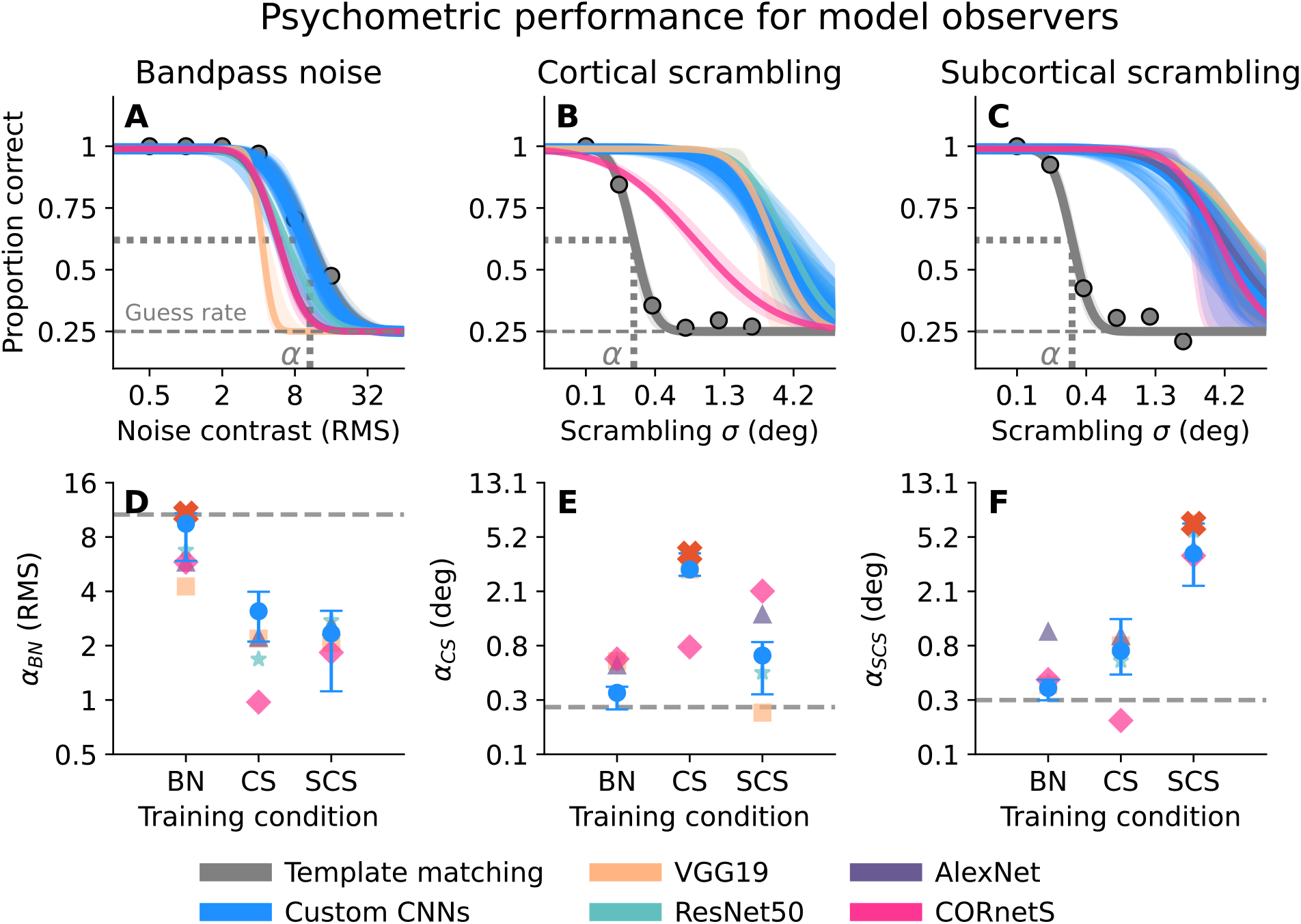
Performance of models in the letter identification task. The top row shows the psychometric functions for models performing the task in the bandpass noise (BN) (**A**), cortical scrambling (CS) (**B**), and subcortical scrambling (SCS) (**C**) conditions. Twenty CNNs had custom architectures randomly generated from an architecture search. Transfer learning was done on four pre-trained networks: VGG19, AlexNet, ResNet50, CORnetS. Shaded region shows 95% CI obtained from 200 bootstrap fits. The bottom row (**D-F**) shows threshold *α* for all CNN models. Threshold *α* is defined as the point on the psychometric function that corresponds to 62% correct (shown for TM model in the top row). Dotted grey line shows the *α* of TM model. Blue marker with error bars denotes median with 95% CI for the custom trained CNNs. Orange cross shows the *α* of the best-performing CNN of all tested models. For each noise condition, it was a custom trained CNN. AlexNet trained and tested on CS is removed due to large fitting error of the psychometric function.

**Fig 6.**
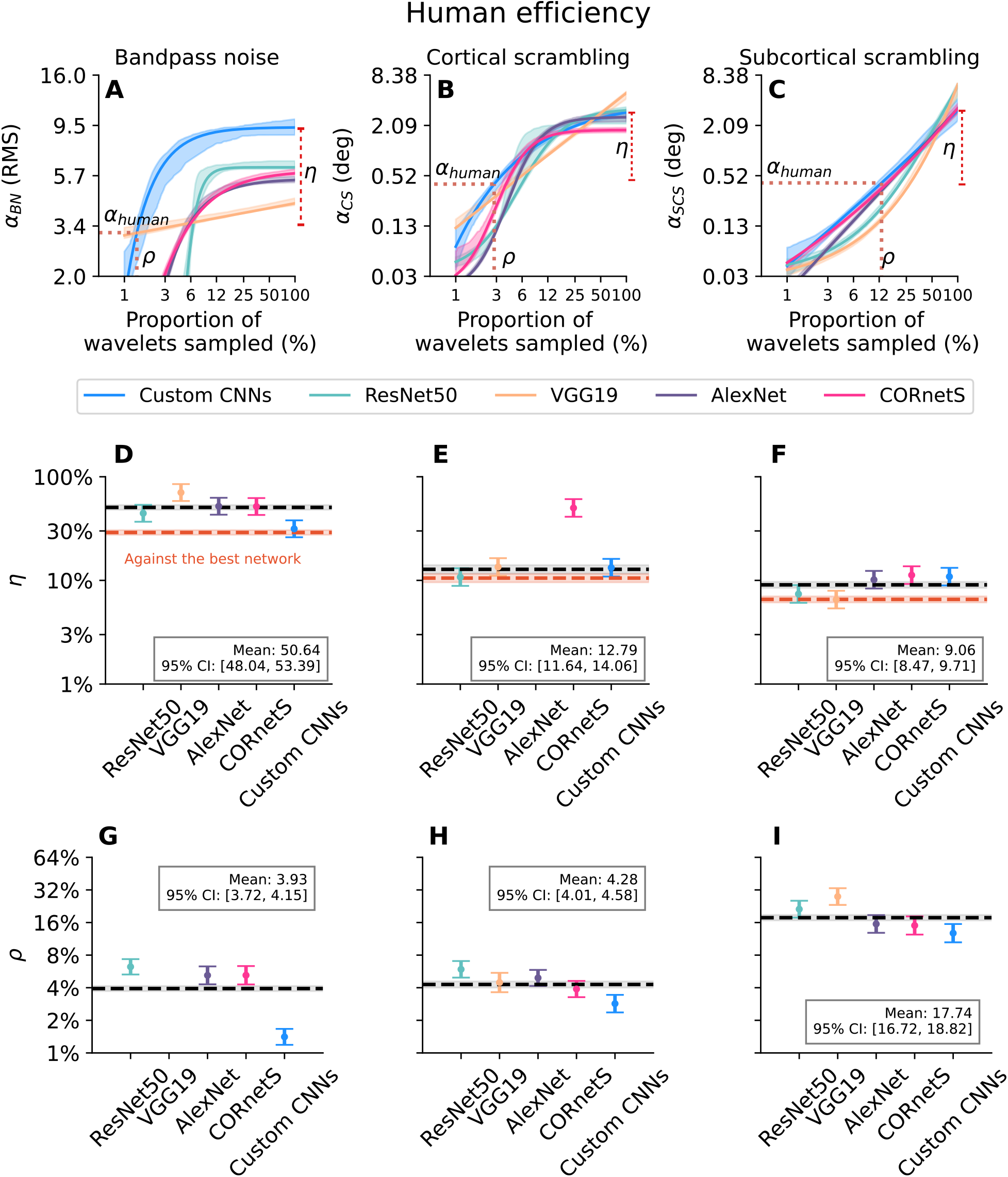
Top row shows threshold *α* of CNNs sampling at different percentages of wavelets in the stimulus for (**A**) BN, (**B**) CS, and (**C**) SCS. Sigmoid or linear function of best fit is shown with 68% CI obtained from 100 bootstrapped fits. Relative efficiency *η* (ranging from 0% to 100%) is defined as the ratio of human *α*_*human*_ to CNN *α* sampling at 100% (**D**-**F**). Sampling efficiency *ρ* is the percentage of wavelet sampled by the CNNs to match *α*_*human*_ (**G**-**I**). For the model *α* of custom CNNs, the mean *α* of twenty CNNs was used. Markers indicate the mean with 95% CI for eighteen subjects across both eyes (two participants removed after outlier removal). Dotted black line denotes the mean efficiency across CNN models with shaded region indicating 95% CI, values of which are shown in the labeled box. Dotted orange line with shaded region denotes the mean *η* compared to the best performing CNN model with 95% CI. The fitting error for AlexNet in CS at 100% wavelets was large and *η* is not shown (**E**). For VGG19 in BN (**G**), there were not enough valid thresholds below human performance for *ρ* to be extracted and is not shown. All function fits with data points are shown in **Fig 5** of Supporting Information.

For the BN condition, we found custom CNN thresholds are not significantly different from TM threshold (Mann-Whitney U test, U=19, p = 0.19). This suggests our architecture search and training procedure produced CNNs that approximated the ideal observer performance for the BN stimulus condition. For the CS and SCS conditions we cannot know how close our custom CNNs are to the theoretical performance limit. In comparisons against previously-published CNNs, we do find that the performance of our custom CNNs reaches or exceeds that of the pre-trained models.

### 2.4 Comparing human performance against that of CNN models

In **Fig 6D-F**, we show the relative efficiency *η* calculated between the human data and the different CNN models. A higher relative efficiency indicates human performance is closer to the performance of the CNNs. The dashed black line shows the mean efficiency across models. For BN, the efficiency was 51%. This means that the human thresholds were approximately half that of the CNN models. For CS stimuli, the efficiency was 13% (AlexNet and CORnetS models excluded as outliers) and for SCS it was 9%. Marginalizing over models, a one-way repeated measures ANOVA was conducted to examine the effect of noise condition on relative efficiency in eighteen subjects. The analysis revealed a statistically significant main effect of noise condition (*F*_(1.61,27.32)_= 879.68, p < 0.001, Greenhouse-Geisser corrected). Post-hoc pairwise comparisons revealed a significant difference between BN and CS (*t*_17_ = 27.35, p < 0.001), BN and SCS (*t*_17_ = 54.95, p < 0.001), and CS and SCS (*t*_17_ = 7.45, p < 0.001). Compared to the best model (dotted orange line), efficiency was 29% for BN, 10% for CS, and 7% for SCS. One-way repeated measures ANOVA also revealed a statistically significant main effect of noise condition (*F*_(1.61,27.32)_= 612.94, p < 0.001, Greenhouse-Geisser corrected). Post-hoc pairwise comparisons also revealed a significant difference between BN and CS (*t*_17_ = 20.15, p < 0.001), BN and SCS (*t*_17_ = 47.50, p < 0.001), and CS and SCS (*t*_17_ = 10.23, p < 0.001). These results show that the efficiency with which humans identify scrambled letters, when expressed as the magnitude of scrambling required to degrade performance to the threshold level, was significantly higher in CS than in SCS.

### 2.5 Modelling inefficiency as a loss of samples

The human participants in our study never out-performed our model observers (*η* was always *<*100%), demonstrating our CNNs made better use of the information available in the stimulus to perform the task. One possibility is that the humans use an inferior algorithm that leads to information loss. Previous studies have modelled this as the effective number of samples that are being used if the strategy applied to them is optimal (Dakin, 2001; Pelli and Farell, 1999; Dakin et al., 2005). Our wavelet-based stimulus generation algorithm allows us to explore this, with the numerical findings of such an analysis giving insight into the differences between human and CNN performance.

We simulated a loss in the sampling of the wavelets forming our letter stimuli by generating variants that were composed of a reduced number of wavelets (randomly sampled). These were then fed into our CNN models, allowing us to find the degree of under-sampling where CNN performance fell to the human level. This gave an indirect measure of the proportion of stimulus information humans effectively used. The modelling results are shown in **Fig 6A-C**. For all models and noise conditions except for one (VGG19 in BN), the CNN *α* monotonically decreased below *α*_*human*_ as we decreased the number of wavelets in the stimulus (visualization of stimuli).

We fit a four-parameter (slope, vertical shift, horizontal shift, and amplitude) sigmoid function or a simple linear model to the data depending on which model gave lower scores of Bayesian Information Criterion (BIC) (Schwarz, 1978) and Akaike Information Criterion (AIC) (Akaike, 1998). We then used the fit relation to find *ρ*, the proportion of wavelets sampled that would make CNNs have the same threshold as *α*_*human*_. *ρ* for different CNN models are shown in **Fig 6D-F**. A one-way repeated measures ANOVA revealed a significant main effect of noise condition, (*F*_1.76,29.91_ = 1135.78, p < 0.001, Greenhouse-Geisser corrected). Post-hoc pairwise comparisons revealed that BN and CS conditions did not differ significantly (*t*_17_ = -2.11, p = 0.15), while both BN and CS differed significantly from SCS (BN vs.SCS: *t*_17_ = -50.89, p < 0.001, CS vs. SCS: *t*_17_ = -40.61, p < 0.001).

It is interesting to compare the two types of efficiency: *η* and *ρ*. In **Fig 6D-F**, *η* is quantified as the ratio between the thresholds of the humans and the CNN models. This measures how resilient humans were in their ability to identify letters under the different types of noise or scrambling. By that analysis, humans are closer to the CNN performance for BN (50%) than they are for the other noise types, with SCS being the lowest (9%). For *ρ* shown in **Fig 6G-I**, we were instead asking how much information (in terms of the proportion of wavelets sampled) needed to be removed from the CNN’s input before its performance fell to human levels. This change in approach reversed our findings. For BN stimuli, CNNs reach human performance with only 4% of wavelets (i.e. removing 96% of the wavelets from the input). This is the lowest efficiency we find, though for CS it was only a little higher. For SCS, the performance of the CNNs dropped to the human level at 18%. This indicates that the identification of SCS letters is far more dependent on the number of samples than for CS stimuli (as is also evident from the slope of the decline).

## 3 Discussion

### 3.1 Identification of scrambled letters by humans and CNNs

In this study we investigated the impact of physiologically-inspired scrambling distortion on letter identification. We employed an algorithm through which stimuli could be generated that exhibited two possible types of scrambling. These were “cortical scrambling” (CS) where the encoded positions of oriented features are randomly perturbed, and “subcortical scrambling” (SCS) where the wavelets used to reconstruct the image are themselves perturbed. Although the distinction between these two types of scrambling is inspired by different possible sites in a back-pocket model of the visual hierarchy (either at the output from simple cells in V1, or at the input forming those simple cell receptive fields) we do not draw any direct conclusions about the physiological nature of scrambling from this behavioural study. Instead, our results demonstrate these two manipulations can have distinct perceptual impact. CS and SCS degrade different information in the image because networks trained on one type of stimuli performed not as well when faced with the other. We further showed difference in CS and SCS can affect perceived scrambling magnitude in a noise matching experiment as well as give rise to different efficiencies for letter identification.

We leveraged recent advances in deep learning to train CNN models to substitute for the role an optimal “Ideal Observer” would play in conventional noise masking studies to extract efficiency (Yamins and DiCarlo, 2016; Kanwisher et al., 2023; Lindsay, 2021). Our approach relies on training the CNNs to exploit the information in our letter stimuli. We trained both custom models (selected in an architecture search then trained from scratch) and four “off-the-shelf” pre-trained networks. We validated our approach (in part) using a condition where the identification of undistorted letters was made difficult by embedding them in spatially bandpass noise (BN), for which the ideal observer strategy is known (template matching) (Van Trees, 1968; Pelli et al., 2006; Ziskind et al., 2014). Our custom CNNs achieving similar levels of performance as template matching in BN is a sign that the training procedure is optimized for the task for the variety of architectures we used.

Combining our psychophysical data with predictions from the CNN models allowed us to investigate the different task demands and relative efficiency for the three stimulus conditions (BN, CS, and SCS). We quantified human performance relative to that of the CNNs, and present this as our first measure of efficiency *η*. This efficiency measured in bandpass noise (51% across all models; 29% comparing to the best model) was significantly higher than the 10% measured in a similar task by Pelli et al. (2006), who instead used spatially broadband noise. We suggest this difference may be due to our use of bandpass stimuli (reducing cross-channel suppressive interactions) and/or our use of a smaller set of letter identities to be discriminated (4 in our study vs. 26 in Pelli et al., 2006). Comparing the two scrambling manipulations, we find a small but significant advantage for CS stimuli (13% vs. 9% across all models; 10% vs. 7% comparing to the best model). This means that, relative to the CNNs, humans are more tolerant to random translations of the position of oriented features than they are to distortions of the wavelets representing those features.

One limitation of the above approach for our scrambling conditions is that the CNN thresholds are around ten times higher than the human thresholds. This places them so far beyond the human limit for recognising “a letter” (see right column of **Fig 2**) that the computation involved may be quite different from that involved in identifying letters near the human threshold (for example, it may rely more on global statistics than local structure). For that purpose, we performed a second analysis to calculate an alternate measure of efficiency *ρ*. Instead of asking how much more noise or scrambling the CNN could tolerate than a human participant, we instead set the human threshold as our benchmark and asked how much we needed to disadvantage the CNN (by randomly removing samples from its input) before its performance was degraded to the human level. This produced an entirely different pattern of results. We find that humans are remarkably efficient at processing SCS letters: CNNs need 18% of the wavelets to be preserved if they are to reach the human performance level. In comparison, CNNs need only around 4% of the stimulus wavelets to match human performance for BN or CS stimuli. Visualising the stimuli with different noise or scrambling magnitudes and with different proportions of wavelets preserved (**Fig 6-8** in the Supporting Information) gives the subjective impression that the SCS stimuli tend to remain more “letter-like” whereas the BN stimuli are reduced to parts of a letter and the CS stimuli appear as a scattering of wavelets.

Inspecting the top row of **Fig 6**, it seems that our three stimulus conditions may differ in how much redundancy exists in the input. For SCS, any loss of wavelets causes an immediate decline in performance for any of our CNN models. For the other two stimulus conditions we most often see a plateau as the proportion of wavelets declines from 100%, followed by a decline once the proportion drops below some critical value. The only exception is VGG 19, for which we saw immediate decline in performance once wavelets were reduced. We suspect this is due to a defining feature of the network that is it only contains 3 by 3 small filters. Despite transfer learning, the pre-trained weights of the network might still bias it to fully exploit the redundancy in our stimuli. Investigating the redundancy in our stimuli, and whether similar plateaus exist in human performance as a function of the proportion of wavelets preserved, is a topic for future study.

### 3.2 Error similarity between humans and CNNs

Despite the presumed similarity in the fundamental architecture, human and CNNs can be shown to behave differently (Wichmann and Geirhos, 2023). For example, CNN models can be worse than simpler models of early visual processing in predicting the visibility of distortions (Berardino et al., 2017). Previous works have also shown a divergence in the pattern of classification errors between CNNs and humans (Dodge and Karam, 2017; Borji and Itti, 2014; Geirhos et al., 2020; Rajalingham et al., 2018). To investigate this, we carried out a supplementary analysis examining error pattern similarity between our CNNs and humans (**Fig 1-3** of Supporting Information). We normalised our data so that this analysis would be independent of the overall performance of the participant (and so not confounded with the main results). We found errors are more similar among CNNs than humans, reinforcing previous findings (Geirhos et al., 2020). We investigated whether our reduced sampling procedure (that lowered CNN performance to the human level) would also bring the pattern of errors into alignment between CNNs and humans. We found only partial evidence, suggesting this effect for SCS but not for CS. Therefore, even though this manipulation made the CNNs more similar to humans when considering the threshold stimulus magnitude, there remains a gap in fully modelling human responses using CNNs.

### 3.3 Internal scrambling and positional uncertainty in human spatial vision

In noise masking studies, it is typically assumed that the dimensions of the external noise applied to the stimulus correspond to the task-relevant internal noise in the visual system. This is necessary if the main effect of that noise is to increase response variability rather than suppressing the response to a target stimulus through nonlinear interactions (Baldwin et al., 2016; Baker and Meese, 2012). Previous noise masking studies have investigated scrambling or positional uncertainty using relatively simple stimuli synthesised from oriented wavelets (Hess et al., 1997; Baldwin et al., 2017). Others measuring Just-Noticeable-Difference (JND) as a function of pedestal position noise level have used regularly spaced lattice (Morgan et al., 2012) and natural images (Christensen et al., 2019). In all cases, they demonstrated internal sensory thresholds for the coding of position in human spatial vision.

Compared to stimuli used in previous works, our study differs in two major aspects. First, we applied our manipulations to letter stimuli. Letters are moderately complex and understanding how visual system performs letter identification informs us processes underlying recognition of more complex classes of objects (Grainger et al., 2008). While Levi et al. (1997, 1999) used a tumbling letter E, their task was an orientation discrimination task with the stimulus consisted of only 17 Gabors arranged with equal spacing. This is different from our stimulus generation scheme where the letters contain many more wavelets whose position, orientation, and phase are more consistent with internal representation based on the activities of oriented feature detectors in the cortex (see stimulus generation in 4.3). Second, we also investigated the effects of a positional noise affecting the subunits from which oriented receptive fields are formed in our subcortical scrambling noise (SCS). This approach is inspired by projection scatter measured in physiology studies (Tao et al., 2012; Nishimoto et al., 2006). In Section 4.3, we show using simulations that the effect of our SCS noise is primarily spreading energy of the Gabor into nearby orientations, increasing the orientation bandwidth while keeping in the spatial frequency passband. This differs from the simple in-place orientation jitter implemented in previous studies (Morgan et al., 2008; Christensen et al., 2015; Dakin, 2001). We do not assert our SCS noise describes a complete and accurate account of the physiological basis of scrambling in the human visual system, however the fact that CS and SCS have discriminable effects suggests they may be used to interrogate distinct aspects of the underlying physiological mechanisms. For example, the advantage we find for the dominant eye in the current study (Fig 4) is present only with SCS stimuli. Under our simplified model of the visual hierarchy, the proposed site of SCS would be just prior to binocular combination. As the processing of SCS stimuli appears to be more vulnerable to a reduction in sampling, we theorise that the dominant eye advantage may indicate a physiological bias (where more mechanisms are responsive to the dominant eye). Different sampling properties have been proposed to account for higher position uncertainty in peripheral and amblyopic vision (Levi and Klein, 1986; Levi et al., 1997, 1987). Studies of the effect of reducing the number of samples available to *human* participants (comparing the dominant and non-dominant eyes) may give further insight into this possibility.

Where our results with SCS stimuli hint at possible differences in projections between dominant and non-dominant eye processing, far greater differences have been argued to exist in amblyopia (Hess et al., 1978; Watt and Hess, 1987; Hess et al., 1990; Hess and Field, 1994; Maruya et al., 2025).. It has been suggested that the mis-wirings in connections from the amblyopic eye cannot be compensated for, due to the overriding imposition of a more dominant map from the fellow eye (Hess et al., 1990). Physiology and fMRI studies have shown that amblyopic eye receptive fields are larger and more heterogeneous (Tao et al., 2014; Hussain et al., 2015; Clavagnier et al., 2015; Szinte et al., 2024; Li et al., 2007). This disruption of amblyopic eye filters has been proposed to explain the distorted percepts that can occur in amblyopia (Hess et al., 1978; Sireteanu et al., 1993; Barrett et al., 2003; Maruya et al., 2025; Molaei et al., 2025). Recent work by Maruya et al. (2025) showed the amblyopic eye precept can be modeled as having a different filter set in the steerable pyramid. In ongoing work, we have applied our methods to the study of spatial distortions in amblyopia and found amblyopic eye makes different mistakes than the fellow eye in the classification of letters affected by SCS, and this divergence in mistake patterns correlates with the visual acuity difference (Zhu et al., 2025).

In normal vision, it is proposed that information is “pooled” within larger spatial windows at increasing visual field eccentricities. Humans may have access only to summary statistics of the content within each pooled region (Freeman and Simoncelli, 2011; Balas et al., 2009). Past work has validated this hypothesis by generating model-driven perceptual metamers, images that are physically distinct but perceptually invariant (Freeman and Simoncelli, 2011; Broderick et al., 2023). Future work measuring the limit at which images can be distorted while remaining metameric to both a given (scrambled) model and humans will shed light on scrambling or internal position uncertainty in peripheral vision.

## 4 Methods

### 4.1 Participants

We recruited twenty one adult human participants with normal or corrected-to-normal vision. Five participants (aged 23-26, 2 female, 3 male) were tested in Experiment 1 (Section 4.6), in which stimuli were viewed binocularly. Twenty participants (aged 20-29, 10 female, 10 male) took part in Experiments 2 (Sections 4.7) in which they were tested monocularly by having them wear an eye patch. Eye dominance was determined by the participant’s sighting dominance (hole-in-the-card test).All testing was carried out with informed written consent in accordance with the tenets of the Declaration of Helsinki, and with the approval of the Centre for Applied Ethics at the McGill University Health Centre.

### 4.2 Equipment

Stimuli were presented on a gamma-corrected Display++ LCD screen (Cambridge Research Systems, United Kingdom) with a mean luminance of 58 cd/cm^2^. At the viewing distance of 62 cm, there were 31 pixels per degree of visual angle. The experiment software ran on a Dell XPS laptop (Dell Technologies, Inc.) and was written in Python using the PsychoPy library (Peirce, 2007). All experiments were performed indoors under dimmed lighting. All CNN training was conducted using PyTorch (Paszke et al., 2019) on CUDA-enabled GPUs.

### 4.3 Stimulus decomposition and re-synthesis algorithm

We based our stimuli on those used in previous studies of letter identification in humans. We used the Times New Roman font, consistent with previous studies investigating the optimal spatial frequency channel using bandpass-filtered letters (Chung et al., 2002b). The letter height (typographic x-height) was set to 1.5 degrees of visual angle, at which the optimal spatial frequency would be 1.5 cycles/deg (Majaj et al., 2002; Chung et al., 2002b; Solomon and Pelli, 1994). Four lowercase letters were chosen as the stimulus set: **o, m, d**, and **z**; based on prior work showing they can be easily distinguished from each other (Courrieu et al., 2004; Janini et al., 2022).

We generated our scrambled stimuli using a novel decomposition and re-synthesis algorithm, inspired by work on the activation properties of cortical receptive fields (Daugman, 1985; Field, 1987) and the multi-resolution wavelet transform (Würtz, 1995; Fischer et al., 2007). Our goal here was not to reconstruct the original image, but rather to generate stimuli that allow us to investigate the impact of scrambling on performance for the spatial frequency channel of interest (chosen as that which should optimally support letter identification).

Our decomposition and re-synthesis scheme involved a transformation of basis using Cartesian-separable log-Gabor kernels (Meese, 2010; Baker et al., 2022), which have the convenience of being D.C.-balanced in any phase (Field, 1987). Compared to the more common “polar-separable” log-Gabor, the spatial-domain kernels defined by the Cartesian-separable log-Gabor function do not splay outwards. This gives a form that is more consistent with the single-cell physiology of V1 (Jones and Palmer, 1987).

In **Fig 7** we present an overview of the stimulus generation pipeline. This can be divided into two stages: We first decomposed the original image at different orientations and phases using a log-Gabor filter bank. Then, we re-synthesised the image from the output of that first stage. It is during re-synthesis that we applied our simulated scrambling. For the Bandpass Noise (BN) condition, no scrambling was applied. For cortical scrambling (CS) or subcortical scrambling (SCS) conditions we applied scrambling at different stages in the re-synthesis.

**Fig 7.**
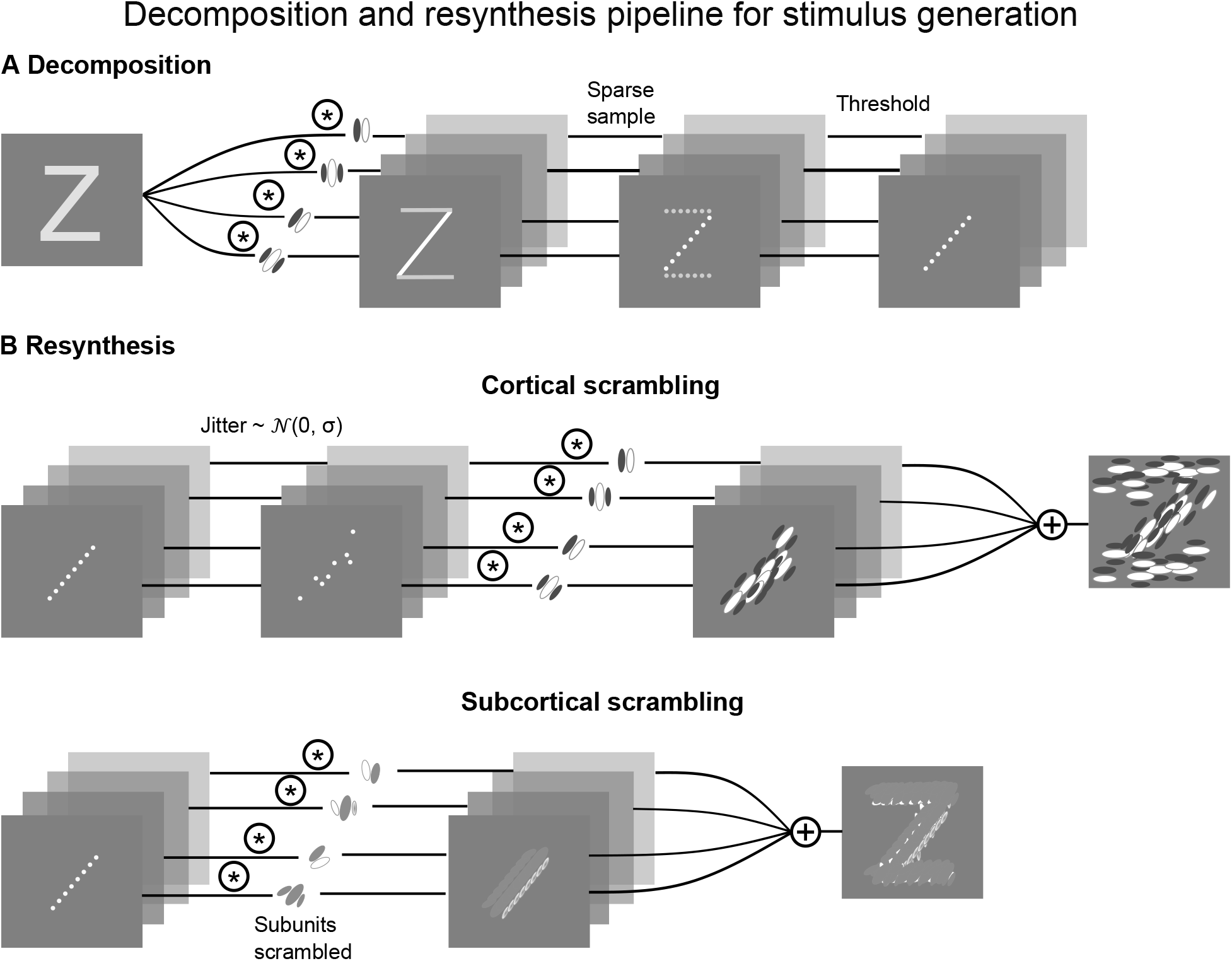
Overview of stimulus generation pipeline. In the decomposition (**A**), the image was decomposed by log-Gabors at different orientations and phases. In the re-synthesis (**B**), the weights were used to synthesize the stimulus made of log-Gabors. For cortical scrambling, we jittered the positions of the weights. For subcortical scrambling, we jittered the positions of the wiring diagram that composed log-Gabor from its isotropic subunits.

In the decomposition stage, we started with raw letter images **X** that were square (200 × 200 pixels). These were filtered by a bank of *m* log-Gabor filters **G**_*j*(*j*∈ Ω)_. Each of Ω = *{*1, …, *m}* indexed a filter defined in Fourier space with a different orientation preference and phase preference. All filters had a peak spatial frequency of 9.8 cycles/image (1.5 cycles/degree) with a bandwidth (full-width at half-magnitude) of 1.5 octaves. There were eight preferred orientations equally spaced between 0° and 360° (orientation bandwidth ±30°). At each orientation, two filters were included in quadrature phase (0 and 90 degrees). This gave a total of *m* = 16 filters. Filtering was performed by multiplication in the Fourier domain

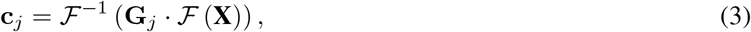

where **c**_*j*_ represented the local weights of a single (j^th^) channel. ℱ and ℱ^−1^ refer to the discrete Fourier and inverse Fourier transform respectively.^2^ Doing this for all the channels gave **c**_*j*(*j*∈Ω)_ which was a stack or “pyramid” of local weights. The decomposition, defined as a wavelet transform, is:

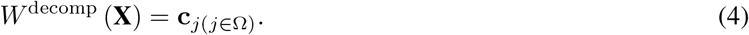

After filtering, we took a sparse sample of each **c**_**j**_ to give a weight value at every half-wavelength of the log-Gabor. We then applied a threshold to retain only the 1% most active weights in the entire stack. This gave the final wavelet weights 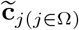.

A letter could then be re-synthesised from these weights using the same filter bank (shifting the 0 degree phase components to 180 degrees, corresponding to complex conjugation in the wavelet transform). For the BN stimuli (without scrambling), we simply convolved 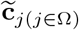 with complex conjugated versions of the filters 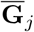 and summed across channels

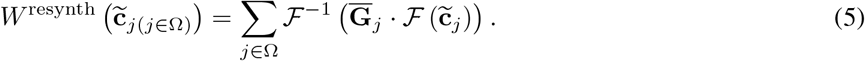

In the bandpass noise (BN) condition, 2D white noise was first fed through the same decomposition and resynthesis pipeline to generate bandpass noise, which was then added to the bandpass filtered undistorted letter.

To simulate cortical scrambling (CS), stimuli were re-synthesised with random position offsets applied to the position of each log-Gabor wavelet. The offsets were drawn from a Gaussian probability distribution, the standard deviation of which controlled the magnitude of scrambling (**Fig 7B, cortical scrambling**). These were applied to the local weights 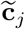, defining the scramble function *S*

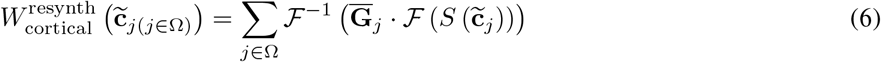

For subcortical scrambling (SCS), scrambling was applied at an earlier stage in the presumed hierarchy by scrambling the log-Gabor wavelets themselves (**Fig 7B**). We first found the wiring diagram that would compose the oriented log-Gabor from isotropic log-Gabor subunits (which have a donut-shaped amplitude spectrum in the 2D Fourier domain) through deconvolution (**Fig 8**). The subunit function was

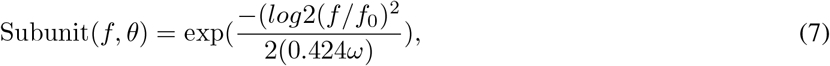

where *f*_0_ is the peak spatial frequency and *ω* is the spatial frequency bandwidth (full-width at half-height) in octaves. For computational efficiency, we kept the top 1% of this wiring diagram and drew position offsets from a Gaussian distribution, the standard deviation of which determined the magnitude of scrambling. The scrambled wiring diagram was convolved with the isotropic subunit to construct a scrambled log-Gabor. This defines the scramble function *S′* which was applied to the filters 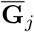 in the SCS re-synthesis

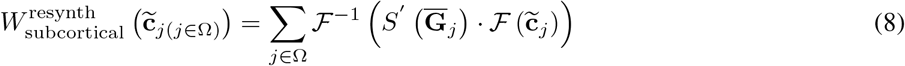

**Fig 8.**
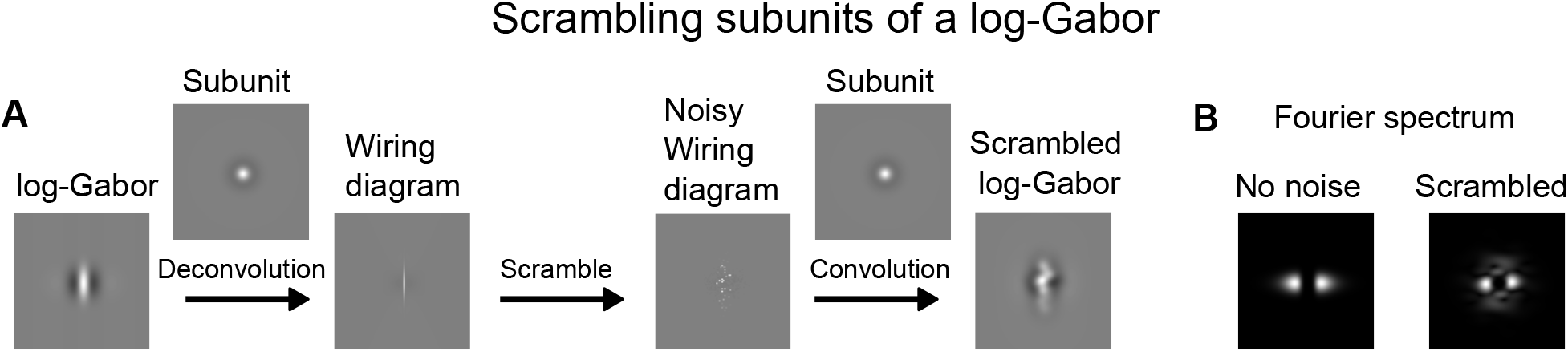
Panel (**A**) shows the generation of a scrambled log-Gabor that was used for subcortical scrambling condition. The log-Gabor was first deconvolved with an isotropic subunit to obtain a wiring diagram. The wiring diagram was then scrambled to simulate miswiring in the subunits. Panel (**B**) shows the amplitude spectrum of the log-Gabor before and after scrambling.

A single scrambled wavelet was generated for each orientation and phase used in the reconstruction for computational efficiency (Maruya et al., 2025). Physiologically however, we would expect the scrambling affecting each receptive field to be distinct.

In **Fig 9**, we summarise the spatial frequency and orientation tuning functions of SCS log-Gabors at three different scrambling levels (based on 200 simulations per plot). Scrambling had the effect of spreading energy from the original orientation channel into neighbouring channels (increasing orientation bandwidth for higher scrambling levels, with the orientation tuning function becoming flat in the limit). We also note a small downward shift in peak spatial frequency and spatial frequency bandwidth.

**Fig 9.**
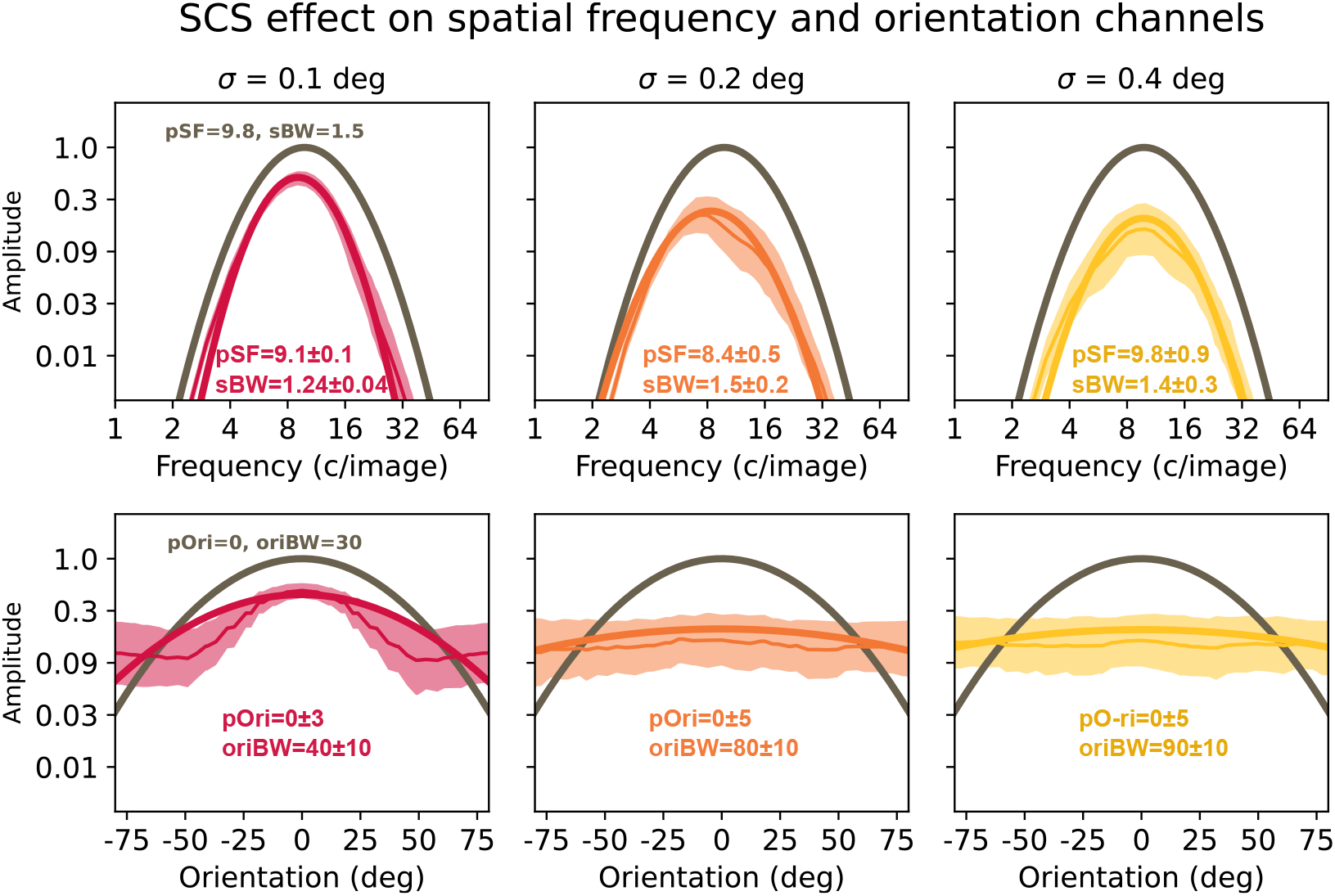
Impact of Subcortical Scrambling (SCS) on the spatial frequency and orientation tuning of log-Gabors. Original tuning functions are shown in grey. Spatial frequency tuning was found by taking a radial slice (in the Fourier domain) of the scrambled log-Gabor (at its optimal orientation before scrambling). Orientation tuning was found by taking a curved slice at the radius equal to the optimal spatial frequency. We scrambled 200 log-Gabors at three scrambling levels and here plot the mean amplitude values with their standard deviations (shaded regions). Four parameters of the tuning functions were fit to each of the 200 samples: peak spatial frequency pSF (cycles/image), spatial frequency bandwidth sBW (octaves), peak orientation pOri (degrees), and orientation bandwidth oriBW (degrees). We report the mean and standard error of the parameters and plot the tuning functions that they would give rise to (coloured curves).

### 4.4 Psychometric function fitting

For baseline contrast sensitivity experiment (Sections 4.5 and 4.7), threshold stimulus magnitudes were found by fitting cumulative normal psychometric functions. For each function, we fit the threshold (for 62% percent correct) and psychometric slope as free parameters. The guess rate and lapse rate were fixed at 25% and 1% respectively. Fitting was performed through a Maximum Likelihood fitting procedure (after Kingdom and Prins, 2016) using the `scipy.optimize.minimize` function from Python’s SciPy package (Virtanen et al., 2020). We calculated the standard error of each fitted threshold using non-parametric bootstrapping. We visualized the histogram of fitting errors for both humans and CNNs and removed outliers beyond the outer fence using Tukey’s method (Tukey, 1977).

### 4.5 Baseline contrast sensitivity experiment: Measuring threshold contrast for bandpass letter identification

To account for individual differences in contrast sensitivity, the stimulus contrast in our main experiment (Experiment 2, see Section 4.7) was set to a multiple of the threshold contrast measured in each eye for each participant. To measure these thresholds, we had participants perform a single-interval four-alternative forced-choice letter identification task. The stimuli were our spatially bandpass letters re-synthesised without scrambling.

For a stimulus **L** with local luminance *L*_*i*_ at each of *n* points and mean luminance 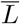, we can calculate its contrast either as the root-mean-square (RMS) value (Moulden et al., 1990; Bex et al., 2009):

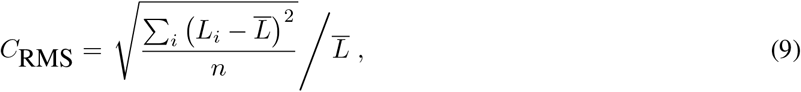

or as the peak Weber contrast (Peli, 1990)

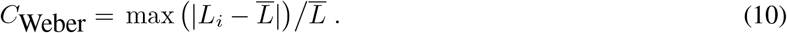

the two differ in that *C*_Weber_ is determined by the contrast of a single point in the image (the greatest deviation from the mean luminance), whereas *C*_RMS_ is proportional to the square root of the contrast energy of the entire stimulus (for a fixed size *n*). They can therefore be thought of as measures of “local” and “global” contrast respectively (Meese et al., 2017). Linear contrast scaling of any specific stimulus image will have equivalent effects on the two contrast measures, though the overall offset between the two will vary on a per-image basis. The stochastic nature of our stimulus generation algorithm for Experiments 1 and 2 could randomly introduce contrast peaks that would have large effects on Weber contrast without making the task easier. On that basis, we chose to measure and control RMS contrast in our experiments. Where convenient, we also report the expected Weber contrast of a stimulus.

The stimuli had a duration of 250 ms above half-magnitude (stimulus contrast controlled by a raised-cosine envelope with 50 ms ramp and 200 ms plateau). The participant identified which of the four letters was shown, and were then given feedback as to whether they were correct. The RMS contrast of the letters was controlled by a two-down one-up staircase (starting level equivalent to 7% Weber contrast, with step sizes set to a factor of 1.6 for the first 4 reversals and 1.3 thereafter). The staircase terminated after a minimum of 30 trials and 8 reversals. The measurement was repeated three times for each eye, with thresholds then obtained through psychometric function fitting (Section 4.4).

### 4.6 Experiment 1: Measuring perceived magnitude of scrambling

We quantified the relative subjective magnitude of the two types of scrambling (CS and SCS) using the matching paradigm. Participants were shown three scrambled letters, positioned in a horizontal row. The letters were spanned 20 degrees of visual angle, with the central letter placed halfway (10 deg) between the left and right letters. All three letters had the same identity (**o, m, d**, or **z**). The central letter was not scrambled: its purpose was to ensure the participant knew the identity of the three letters. The “reference” letter, presented on the left, had a fixed scrambling magnitude from a set of five logarithmically-spaced levels. The “match” letter, presented on the right, had an adjustable scrambling magnitude. On each trial, the scrambling applied to the match letter was adjusted by the participant until they were satisfied that it appeared “equally scrambled” as the reference.

Four matching conditions were tested, with matches both between the same type of scrambling (matching CS to CS, or SCS to SCS) and between different types of scrambling (matching CS to SCS, and vice-versa). For each of the four letter identities and the five reference levels, the participant performed two matches: one where the match letter began with a “high” level of scrambling and another where its initial scrambling was relatively “low”. This gave 40 trials for each matching condition, which were performed in a randomised order.

To encourage participants to make a holistic judgement of scrambling magnitude, the scrambled letter stimuli in this experiment were dynamic. Both the reference and match letters were updated at 5 Hz, being replaced by another scrambled letter of the same identity and scrambling magnitude. Participants viewed the stimuli binocularly, with stimulus RMS contrast set to ten times the mean contrast threshold of the five participants to ensure visibility. The total range of Weber contrast for all stimuli was 57-72% for CS and 57-70% for SCS.

### 4.7 Experiment 2: Measuring noise threshold for letter identification

Similar to the measurements of contrast threshold in baseline contrast sensitivity experiment (Section 6.2), our main experiment was a single-interval four-alternative forced-choice letter identification task. In this experiment though, the contrast of the letters was fixed at an RMS contrast four-times the threshold contrast measured in each eye for each participant. Converted to Weber contrast, the mean contrasts used were 5% for BN, 11% for CS, and 10% for SCS (for both the dominant and non-dominant eyes). We instead varied the magnitude of the noise or scrambling applied to the letter to find the threshold level where performance reduced to threshold (62% percent correct). This requires a flipped staircase, as increasing the contrast of the bandpass noise in the BN task, or the scrambling magnitude in the CS or SCS task made the task more difficult (where increasing contrast in the previous task made it easier). A two-up one-down staircase controlled the noise level of the stimulus, with the same terminating criteria as used in baseline contrast sensitivity experiment. There were three repeats of each eye and noise condition.

### 4.8 Training and selecting custom CNNs

We generated three sets of stimuli to train the CNNs, one for each noise condition (BN, CS, and SCS). Each comprised of stimuli at six log-spaced magnitudes that covered the entire range used in our human testing. At each magnitude, we generated 50 examples for each target letter. This resulted in a training set of 1200 stimuli. During training, we used a split of 70%, 20%, and 10% for training, validation, and testing. Each batch consisted of 32 images. Stimuli in all datasets were max-min normalized following 8-bit quantization in the grey level (preventing the CNN having access to more information than our human participants).

To optimise the CNN architecture for each condition, we performed an architecture search (**Table 2**). Values were drawn for each hyperparameter from a discrete uniform prior. We randomly sampled 100 architectures from this grid, and selected the top 20 by ranking their validation accuracy after training for 10 epochs. Histograms of hyperparameters for the generated and selected architectures are shown in **Fig 4** of Supporting Information.

The top 20 architectures we chose were then trained from scratch for 100 epochs. Early stopping was used with the patience parameter set to 10. All architectures had the same max pooling kernel size of (2,2), as larger kernel sizes frequently led to invalid architectures. The first and second convolutional layers always had 32 and 64 channels respectively to ensure good feature extraction early in the network. Each convolutional layer was always preceded by batch normalization (apart from the first) and followed by the rectified linear unit (`ReLU`) activation function. We used one fully connected layer after the last max pooling layer and before the output layer for which the `SoftMax` activation function was used. We used the `SparseCategoricalCrossentropy` objective loss function (Chollet, 2021) and the Adam optimizer (Kingma and Ba, 2014) to minimize this loss function with a fixed learning rate of 0.001. All model predictions presented in this paper were made on a a test dataset with 1200 stimuli (50 stimuli per letter and noise level for four letters and six noise levels).

### 4.9 Transfer learning using pre-trained networks

We employed transfer learning with four convolutional neural network architectures: ResNet-50, AlexNet, VGG-19, and CORnet-S pre-trained on ImageNet (Krizhevsky et al., 2012; He et al., 2015; Simonyan and Zisserman, 2015). To accommodate single-channel grayscale input, we modified the first convolutional layer of each network by averaging the pre-trained RGB weights across color channels, preserving the learned edge and texture features while adapting to our stimulus format. The final classification layers were replaced with a new fully-connected layers matching our 4-class letter identification task. Training proceeded in two phases: first, we froze the convolutional backbone and trained only the modified first convolutional layer and new classification head for a minimum of 10 epochs using the Adam optimizer again (Kingma and Ba, 2014) with a learning rate of 0.001; second, we unfroze all network parameters and fine-tuned the entire model with a reduced learning rate of 0.0001. We trained for the same number of epochs as custom CNNs (100 epochs) and early stopping patience was also set to 10 epochs monitoring validation loss to prevent overfitting in each phase.

### 4.10 Template matching model

We also implemented a template matching model, which performed an in-place pixel-by-pixel multiplication between each noiseless bandpass letter with the noisy stimulus. The “max” rule was used to pick the letter that gave the highest standard deviation in the response distribution. The predictions of the template matching model were made on the same test set as the CNNs.

## 5 Additional information

### 5.2 Author contributions in CREDIT format

**RXZ**: Conceptualization, Data Curation, Formal Analysis, Funding Acquisition, Investigation, Methodology, Software, Visualization, Writing - Original Draft. **ASB**: Conceptualization, Formal Analysis, Funding Acquisition, Methodology, Project Administration, Software, Supervision, Visualization, Writing - Review & Editing.

### 5.3 Data availability statement

All relevant human and model data, CNN training testing datasets, CNN models can be found on figshare at 10.6084/m9.figshare.25631925 (at publication). All experiment, simulation, analysis, CNN training code can be found on figshare at 10.6084/m9.figshare.25631934 (at publication).

## Acknowledgements

This work was supported by an NSERC Discovery Grant (RGPIN-2022-04216) awarded to ASB, a CIHR Vanier Doctoral Scholarship awarded to RXZ, and the Montreal General Hospital Foundation (through their support of the Centre for Digital Brain Therapies). The authors would like to thank Robert Hess for discussion and feedback, and members of RXZ’s thesis committee: Curtis Baker, Pouya Bashivan, and Christopher Pack for helpful comments and guidance.

## Supporting Information

### 1 Nested model hypothesis testing

Nested model hypothesis test results are shown in **Table 1**.The reduced model is the linear model for the SCS test CS match condition, which is transformed to fit the data for the CS test SCS match condition. The full model refers to fitting two linear models to two conditions separately. We find that the “full” model (where matching SCS to CS is not equivalent to matching CS to SCS) gives a better account of our data. When comparing the difference between these two models, we find that assuming the reduced model would result in a less than 10% difference for the majority of the test range. For the first and second highest scrambling levels tested in the CS test SCS match condition, the difference was 12% and 45% respectively.

**Table 1.** Comparing full model with reduced model for the matching data using nested model hypothesis test.

| $SSE_{full}$ | $SSE_{reduced}$ | $\Delta SSE$ | $\Delta p$ | $MSE_{full}$ | F statistic | p-value |
| --- | --- | --- | --- | --- | --- | --- |
| 0.03 | 0.17 | 0.14 | 2 | 0.005 | 13.91 | 0.0056 |

**Table 2.**
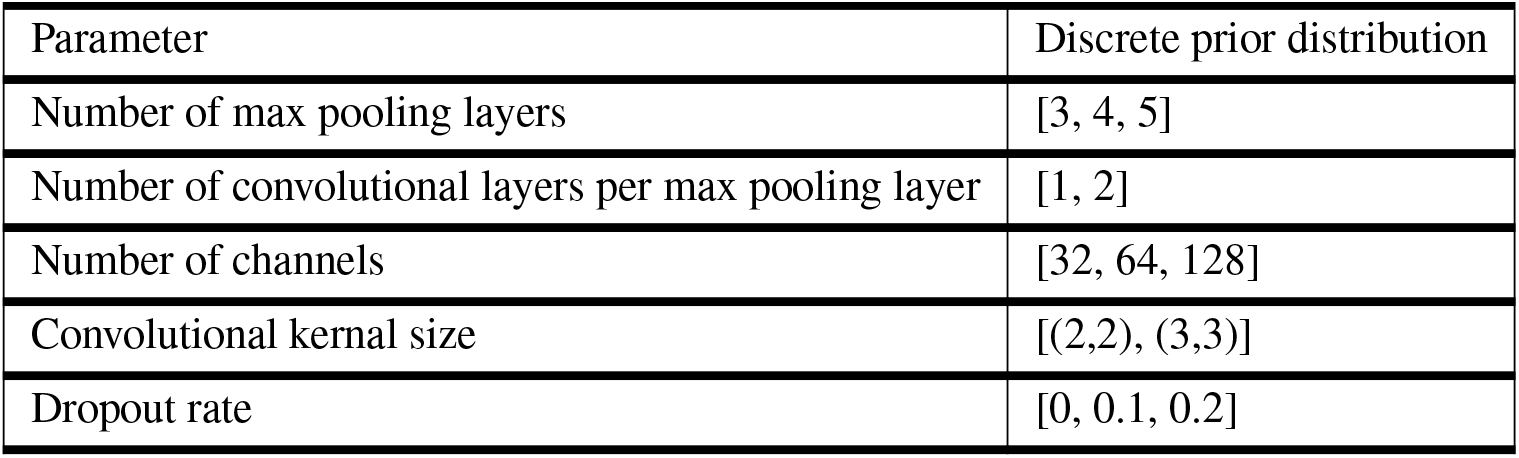
Discrete prior distributions for the architecture search. Values were drawn from a uniform distribution.

### 2 Contrast thresholds for bandpass letter identification

We measured monocular contrast thresholds for identifying letters shown to the Dominant (D) and Non-Dominant (ND) eyes of each participant. The stimuli used in these measurements were re-synthesised (spatially-bandpass) letters without any added noise or scrambling. Across our 20 participants, D eye thresholds (mean *C*RMS 0.0036 95% CI [0.0033 to 0.0039]) did not significantly differ from those in ND eyes (0.0038 95% CI [0.0034 to 0.0041]) when compared using a paired t-test (*t*_19_=-1.08, p=0.29). These RMS contrast thresholds were equivalent to an expected Weber contrast of 2.5% for D eye and 2.6% for ND eye.

### 3 Comparing error patterns

To investigate the error patterns, we first computed confusion matrices for each human and CNN. To ensure the fairness of the comparison between humans and CNNs, we generated another dataset with noise levels that humans were shown in the staircase. For each pairwise comparison, the dissimilarity of errors was calculated separately for each target letter first and added together. For the metric of dissimilarity, We calculated the Kullback–Leibler (KL) divergence between each probability vector and the average of the two probability vectors before suming them. The higher this value the more *dissimilar* the probability distributions are from each other. This metric is symmetric and circumvents the undefined case of any entry in the true distribution being zero while the approximate distribution entry being non-zero. Our metric has the form:

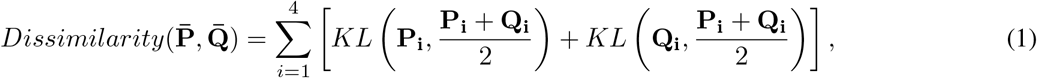

where 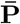 and 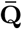 are confusion matrices excluding the diagonal elements, **P**_**i**_ and **Q**_**i**_ are response probability vectors of length 3 for the ith target letter. Our metric is bounded between 0 (same distribution) and 8log(2) (disjoint distribution).

We were interested in whether errors made by CNNs were similar to human errors. Furthermore, we wondered whether constraining CNN performance to human level with reduced sampling could potentially align errors made by humans and models. For reference, we also computed similarity of errors for humans compared to other humans, and CNNs compared to other CNNs. Example dissimilarity matrices are shown in **Fig 1** (100% wavelets kept for CNNs, SCS).

**Fig 1.**
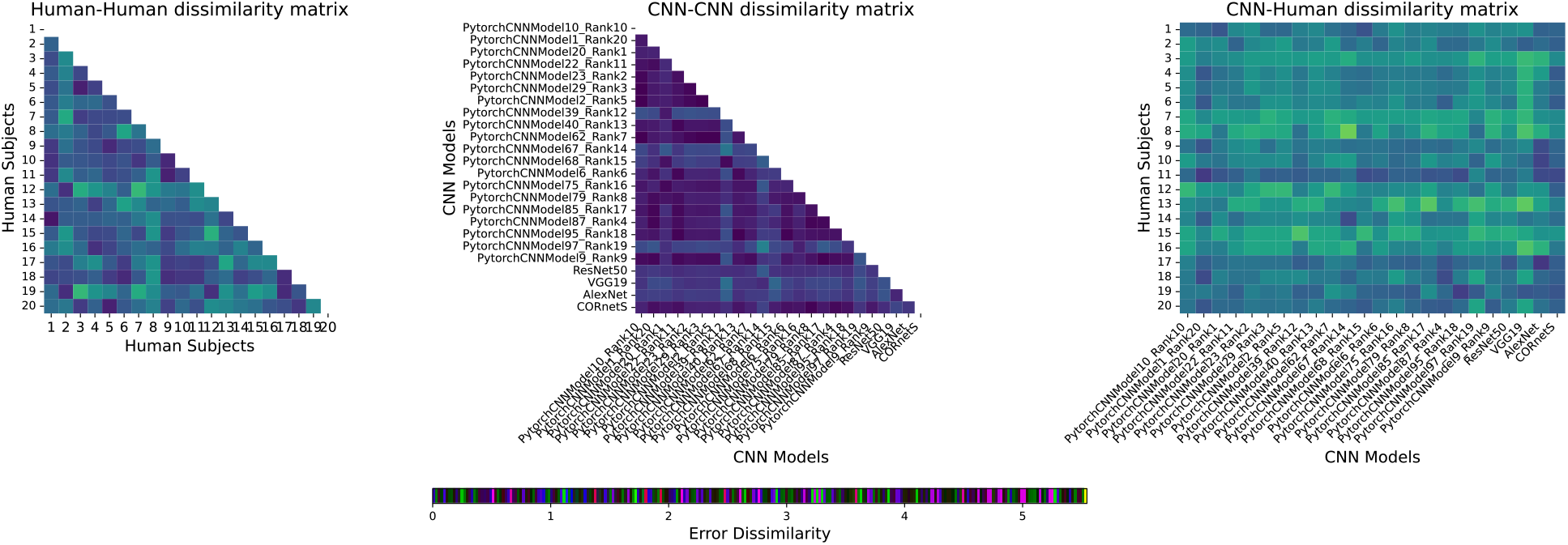
Pairwise dissimilarity for humans against humans, CNN against CNNs, humans against CNNs. Noise condition is SCS. Human data are from dominant eye viewing condition. CNNs are sampling 100% of the wavlets.

The complete results of comparing error patterns are shown in **Fig 2** and **Fig 3** for humans viewing through their dominant eye (DE) and non-dominant eye (NDE), respectively. We found that across proportions of wavelet sampled, CNNs had higher dissimilarity to humans than humans to other humans for CS (Mann-Whitney U test. D: U=1442, p<0.0001; ND: U=1334, p=0.0009) and SCS (D: U=1206, p=0.043; ND: U=1288, p=0.0058). Interestingly we found a significant linear relationship for SCS (D: slope parameter, *t*_6_=3.35, p=0.015, *R*^2^: 0.65; ND: slope parameter, *t*_6_=3.64, p=0.011, *R*^2^: 0.69) where decreasing the percent of wavelet sampled aligned the CNN errors to be closer to humans. For BN, we found CNN errors were not significantly different from human errors (D: U=1045, p=0.87; ND: U=1033, p=1.03). In all cases, we found CNNs were more similar to other CNNs than humans to other humans for both dominant eye (BN: U=53, p<0.0001; CS: U=21, p<0.0001; SCS: U=53, p<0.0001) and non-dominant eye (BN: U=38, p<0.0001; CS: U=58, p<0.0001; SCS: U=67, p<0.0001).

**Fig 2.**
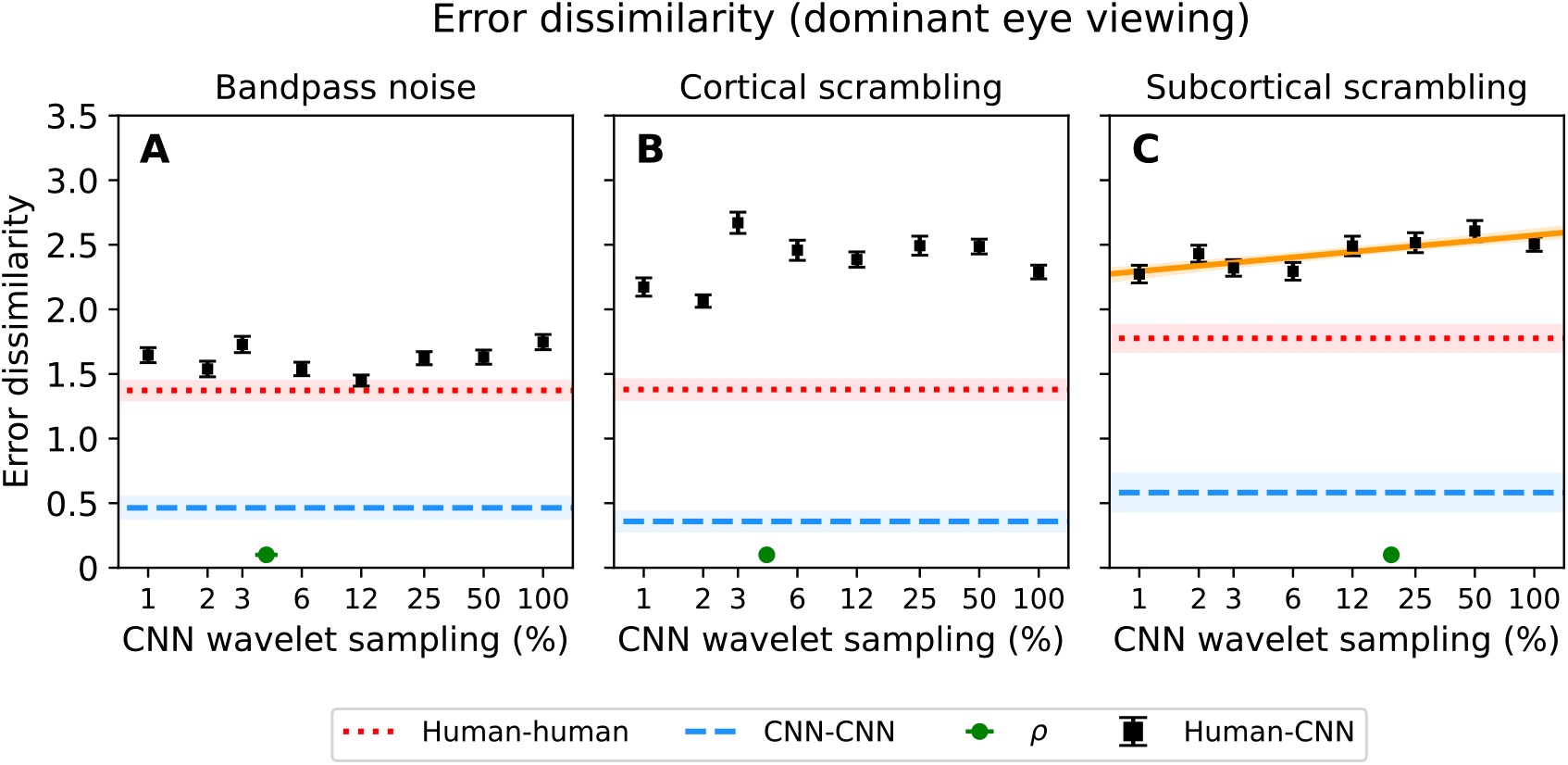
Comparing error dissimilarity between CNNs and humans (dominant eye). Dissimilarity across different proportions of sampled wavelets for (**A** BN), (**B**) CS, and (**C**) SCS. The error dissimilarity metric is based on the Kullback-leibler (KL) divergence. Human to CNN dissimilarity is shown in black markers with mean and 95% CI. SCS is the only noise condition for which the slope is significantly different from zero after fitting a simple linear model. Its line of best fit with 95% CI shaded is shown in orange. For reference, human to human dissimilarity (red) and CNN to CNN dissimilarity (blue) are also shown with 95% CI shaded region. For each noise condition, average *α* across models is shown as a green marker with mean and 95% CI.

**Fig 3.**
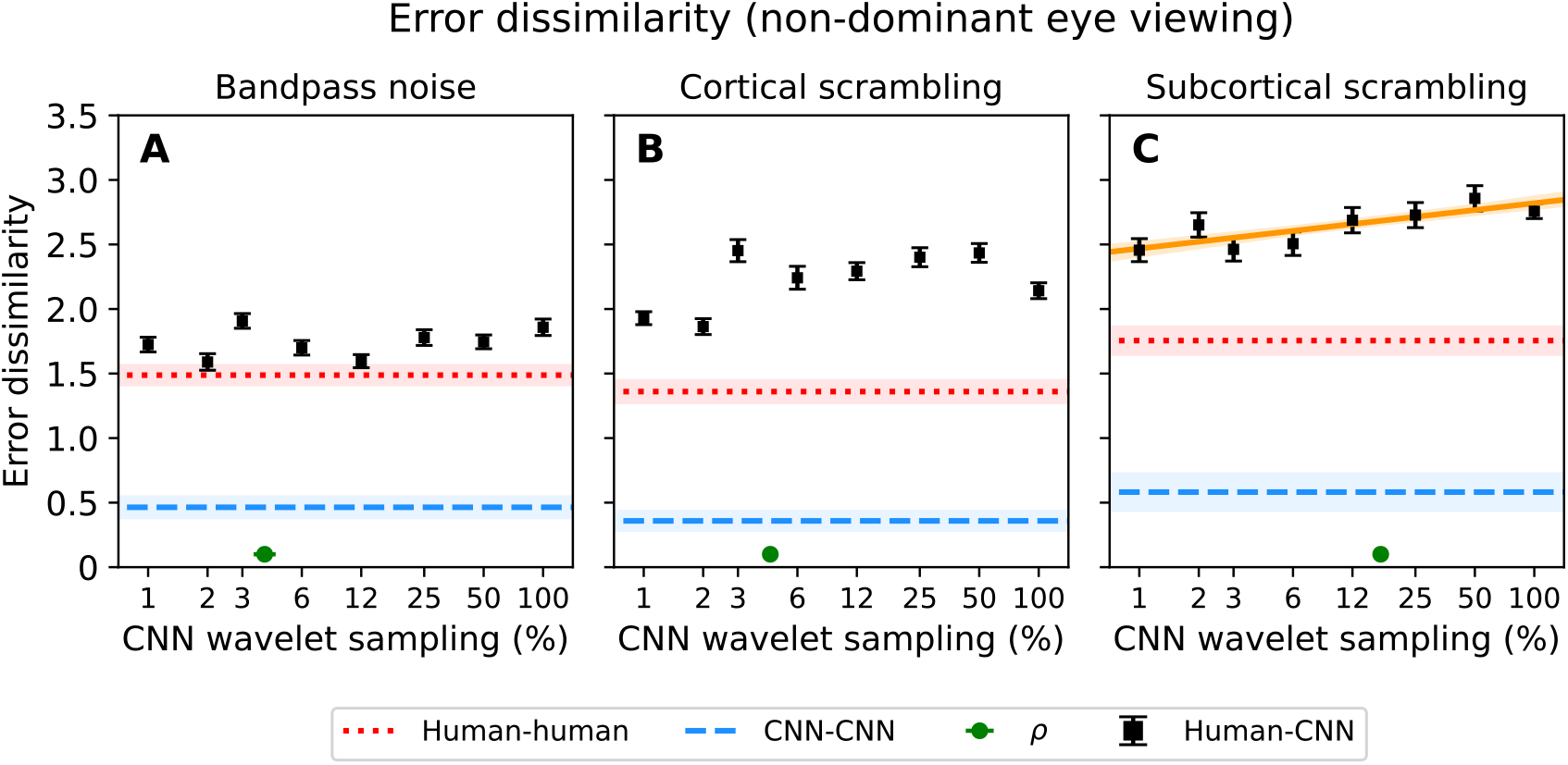
Comparing error dissimilarity between CNNs and humans (non-dominant eye). See captions of **Fig 2** for details. SCS is again the only noise condition for which the slope is significantly different from zero after fitting a simple linear model.

### 4 Architecture search hyperparameter histograms

The architecture search results are shown in **Fig 4**. Out of 100 architectures, top 20 architectures were selected based on validation accuracy after training for 10 epochs. We see chosen architectures tend to have higher numbers of pooling (first row) and convoultional layers (third row).

**Fig 4.**
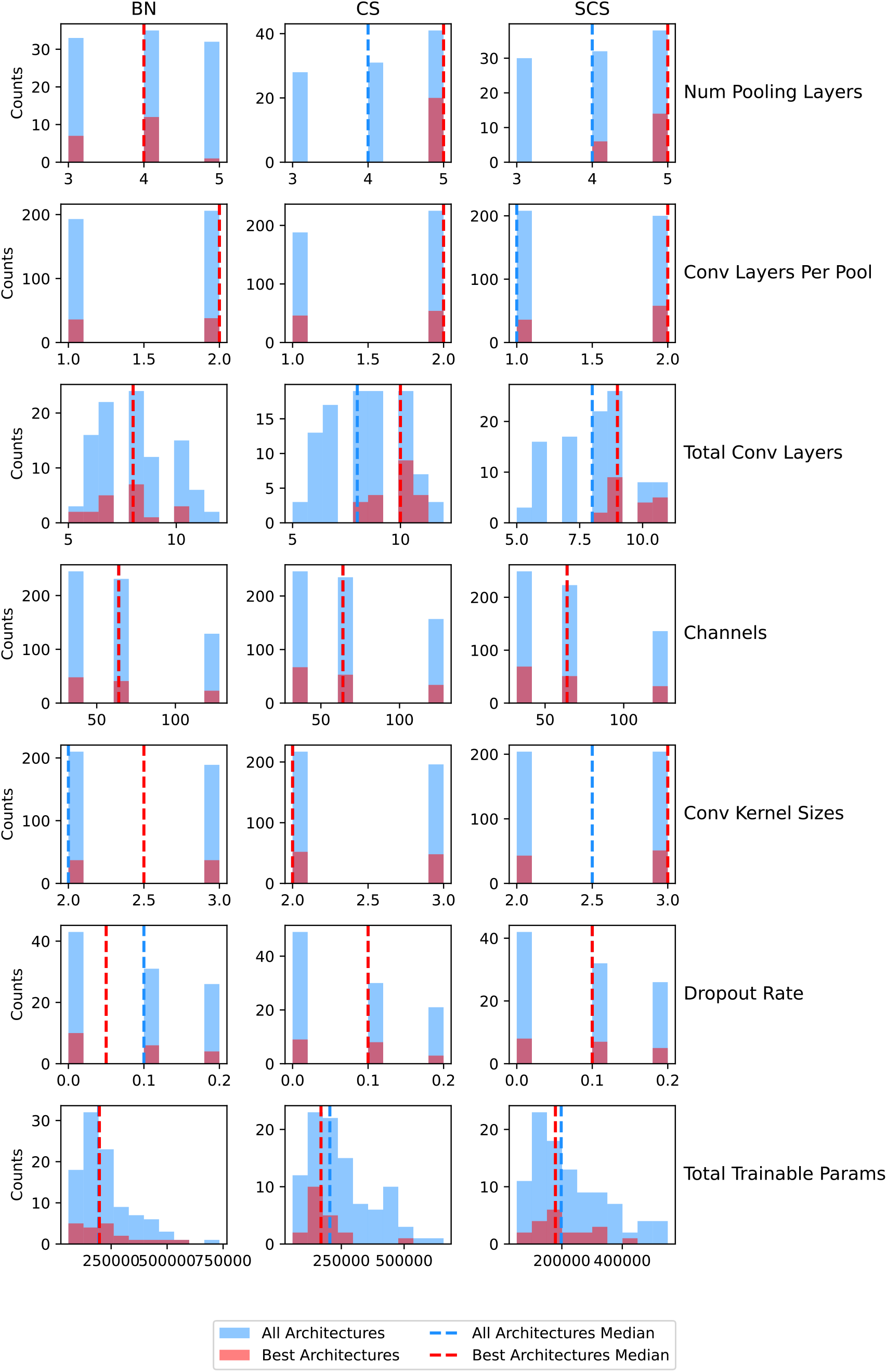
Histograms of hyperparameters of CNNs trained on bandpass noise (BN), cortical scrambling (CS), and subcortical scrambling (SCS) for 100 random architectures searched (blue) and 20 selected architectures (red).

### 5 Individual fits to CNN threshold as function of sampling

Individual function fits are shown in **Fig 5**. CNN performance in bandpass noise and cortical scrambling exhibited a plateau when wavelet sampling was reduced initially and a sharp fall off at lower sampling rate (except for VGG19). In subcortical scrambling, CNN performance decreased immediately once sampling was reduced.

**Fig 5.**
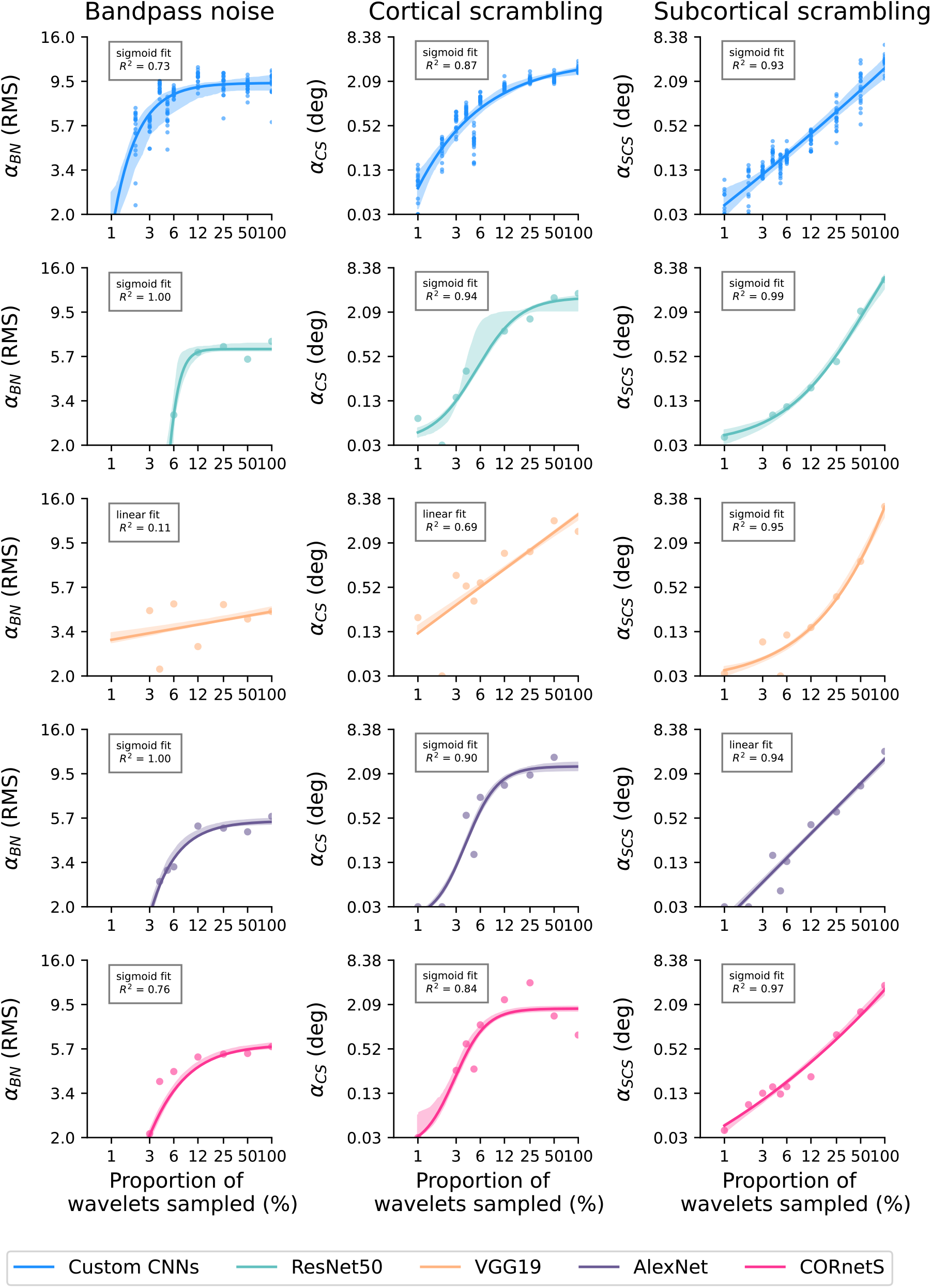
Sigmoid or linear fits to CNNs. Sigmoid model contained four free parameters: slope, vertical shift, horizontal shift, and amplitude. Linear function contained two free parameters: slope and intercept. The fit is shown for the better model defined as the one that gives lower score on Bayesian Information Criterion (BIC) and Akaike Information Criterion (AIC).

### 6 Visualisation of sparse sampling wavelets in the stimulus

We can directly visualize the stimuli at different sampling rate and noise level to qualitatively judge whether what equates CNNs to human performance looks discriminable to humans. The stimuli are inherently stochastic but here we aim to give a high-level impression of what the stimuli look like at various sampling rate and noise level. The visualizations are shown in **Fig 6** to **Fig 8**. If humans were sampling at the same rate ρ that made CNNs have the same performance as humans α they would look like stimuli shown in the second column from the left. For BN and CS, they look like a smattering of wavelets. In SCS, the contour of the letter is more preserved at the cost of losing orientation tuning of individual wavelets.

**Fig 6.**
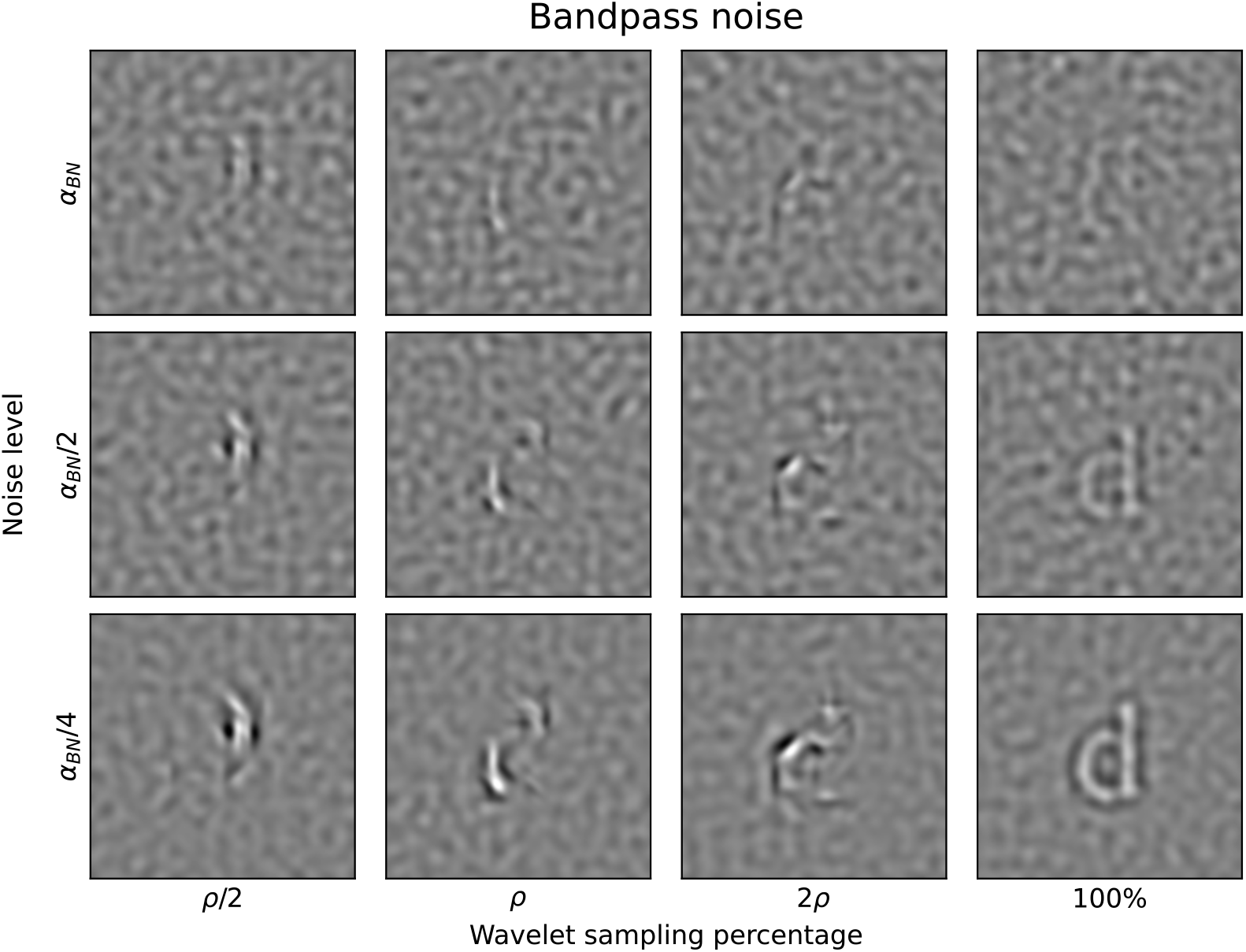
Jointly visualizing the effect of noise magnitude and wavelet sampling on a letter “d” for bandpass noise. *α* indicates the human threshold across dominant and non-dominant eyes. *ρ* indicates the sampling percentage that equated CNNs performance with human performance.

**Fig 7.**
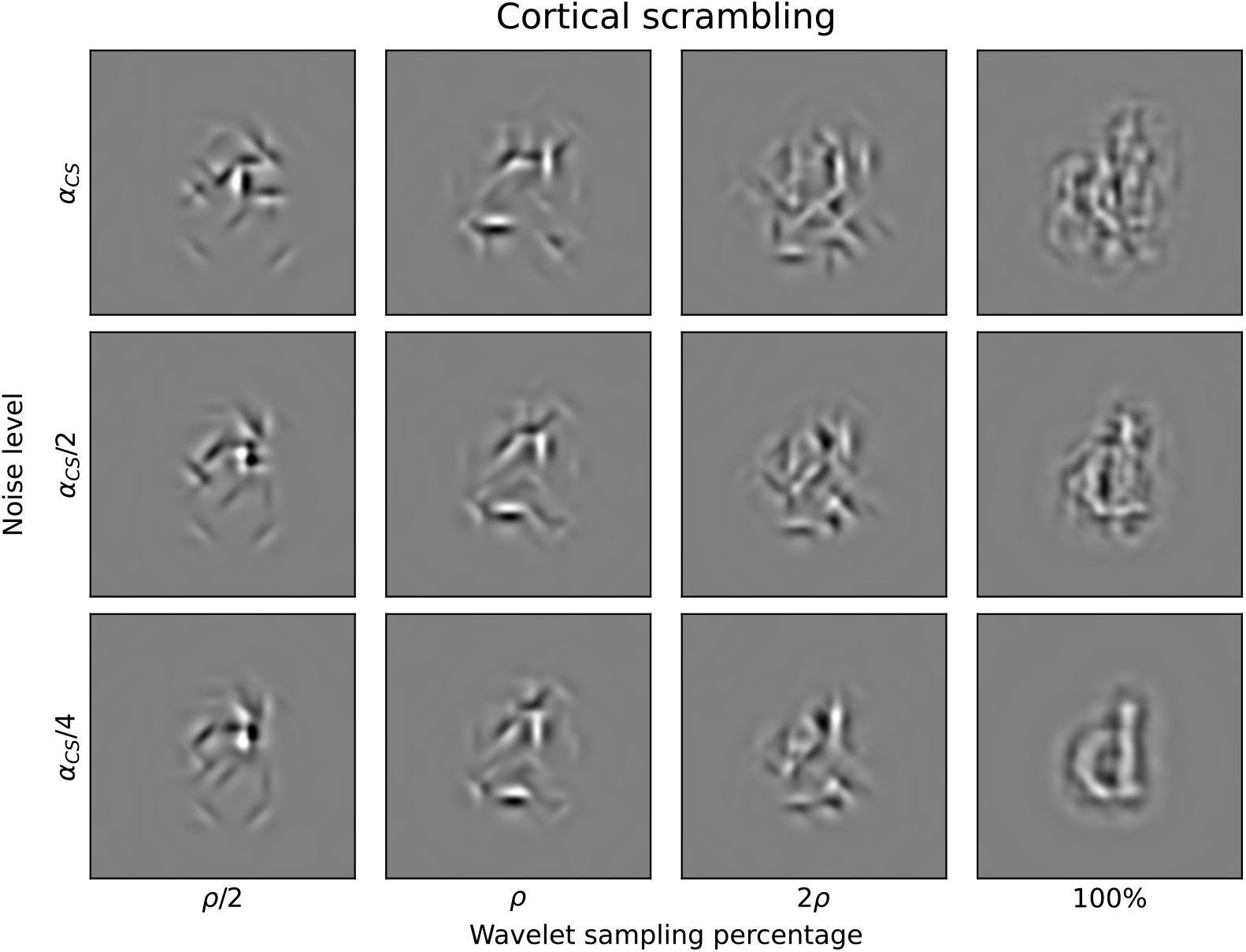
Jointly visualizing the effect of noise magnitude and wavelet sampling for cortical scrambling.

**Fig 8.**
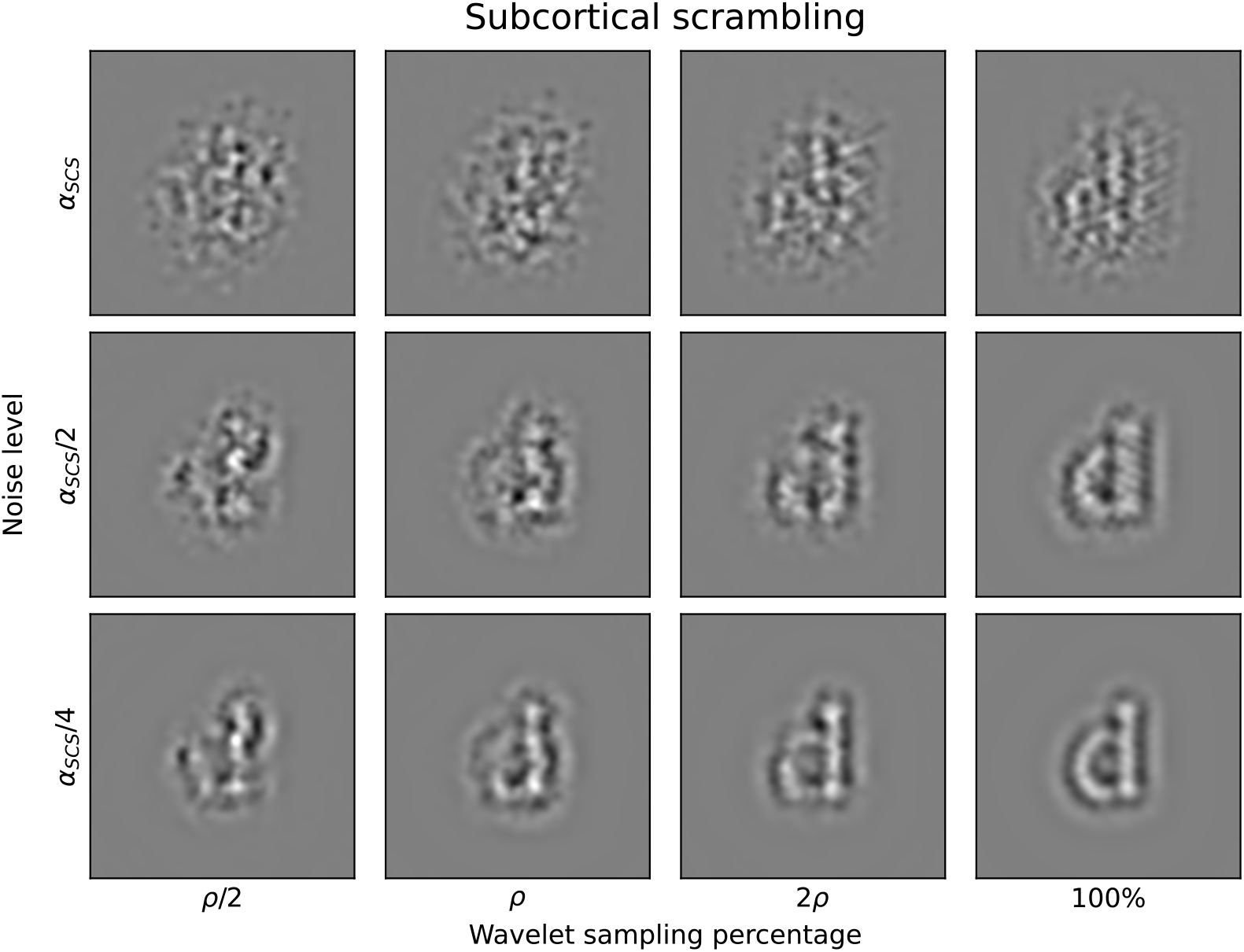
Jointly visualizing the effect of noise magnitude and wavelet sampling for cortical scrambling.

## Footnotes

2 Conversion to and from the Fourier domain was performed using the Fast Fourier Transform module `fft` of the NumPy package in Python (Harris et al., 2020).

